# Coordinated multilaminar dynamics underlie multiplexed computation in macaque motor cortex

**DOI:** 10.1101/2025.11.26.690660

**Authors:** Laura López-Galdo, Simon Nougaret, Demian Battaglia, Bjørg Elisabeth Kilavik

## Abstract

Functional multiplexing is a signature of higher-order brain regions, supported by single-unit mixed selectivity and the recurrent circuitry of the six-layered cortical microcircuit, which allows columns to retain incoming information and integrate it into stable latent representations. It remains unclear whether multiplexing arises from specialized processing within individual layers or from the collective coordination of multiple layers. Here, we analyzed laminar recordings from macaque motor cortex during a delayed match-to-sample task. We identified laminarly distributed, behaviorally specific subspaces that encoded distinct task variables. When the behavioral meaning of a variable was preserved, the same coding subspace was reused across time; when sample information was transformed into a match, the sample subspace was recycled to instead encode stimulus validity. In contrast, identical sensory variables -such as target direction-engaged distinct subspaces when their computational role differed, e.g. during visual encoding versus motor preparation. Subtle variations in laminar weights gave rise to multiple coexistent subspaces, enabling multiplexed representations at the columnar scale. Beyond their coexistence, we tracked the temporal evolution of individual coding subspaces. Task timing paced their geometric reorganization, with task-relevant information propagating along structured trajectories as the laminar center of mass shifted between superficial and deep layers. These laminar trajectories were consistent across sites, though propagation patterns varied, while overall population activity lacked structured dynamics, leaving coherent laminar information to emerge atop a background of unspecific fluctuations.

**Highlights:**

- Coordinated multilaminar activity encodes task-relevant information in motor cortex
- Different layer combinations span multiplexed coding spaces
- Coding spaces for task condition, cue and direction are flexibly repurposed over time
- Task timing paces the geometric reorganization of coding spaces

**Graphical Abstract:** 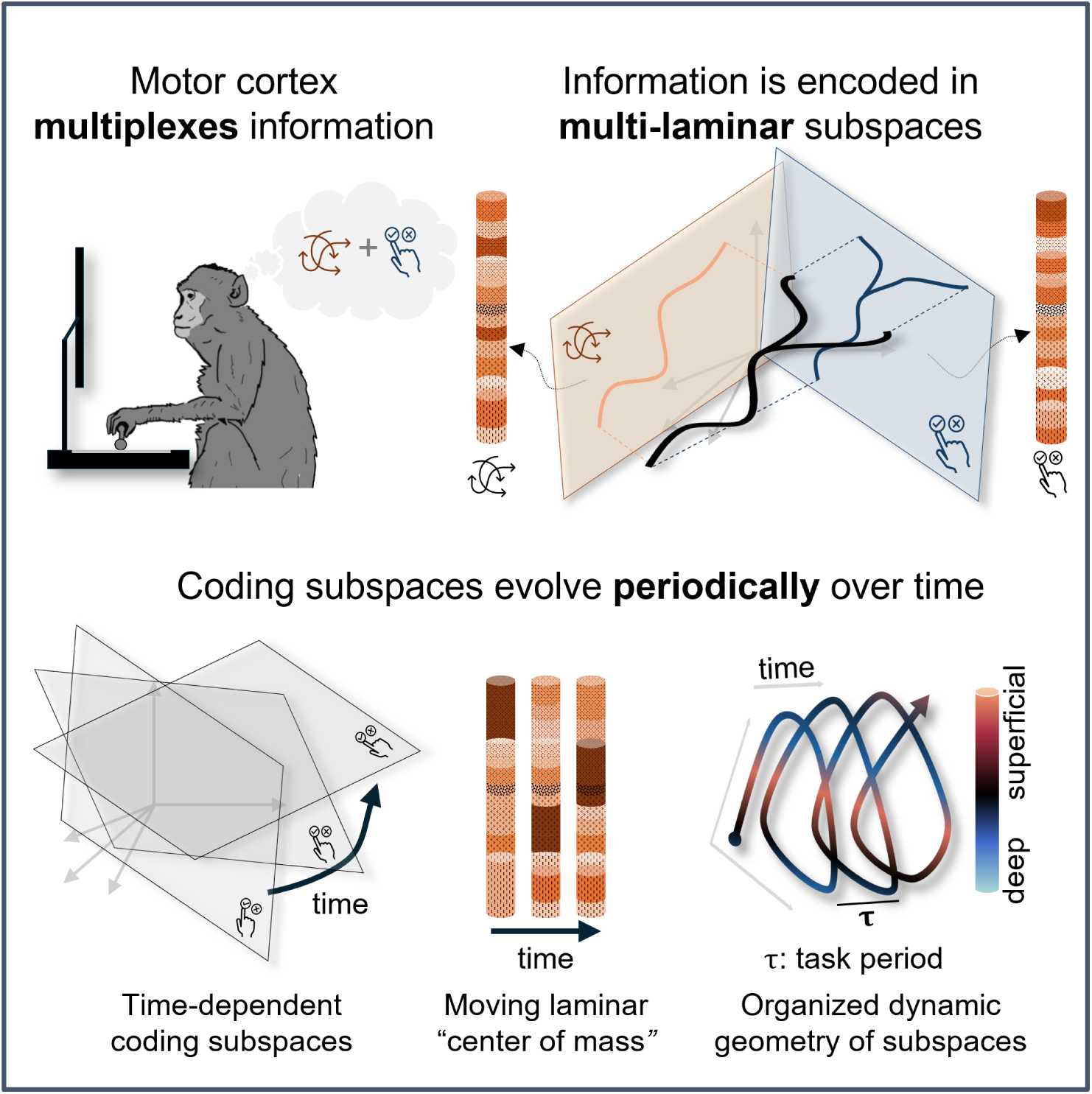

## Introduction

Higher-order cortical areas excel at multiplexing. Their ability to process several kinds of information in parallel relies on the mixed selectivity of single units, the flexible interaction of neural populations, as well as the generation of functional cell assemblies **(Buzsáki, 2010; Rigotti *et al*., 2013; Fusi, Miller and Rigotti, 2016; Tye *et al*., 2024)**. The canonical cortical microcircuit has been proposed as an optimal substrate to perform such computations **(Mountcastle, 1997; Douglas and Martin, 2004; Bastos *et al*., 2012)**. Canonical circuit descriptions model the vertical organization of the six-layered neocortex as an architecture characterized by strong recurrent connections within layers and largely stereotyped patterns of input-output connections between them. This canonic connectivity enables cortical columns to buffer in time multiple incoming transient inputs in parallel, so that the information they convey can be combined into stable latent representations **(Nolan *et al*., 2025)**. The question remains open, however, of whether such multiplexed computation is performed through parallel processes taking place simultaneously in distinct specialized layers **(Bastos *et al*., 2012; Mendoza-Halliday *et al*., 2024)** or, rather, through the collective coordination of multiple layers, operating in an integrated and not easily separable fashion **(Nigam, Pojoga and Dragoi, 2019; Nolan *et al*., 2025)**. This is especially relevant in motor cortex, where laminar organization is pronounced than in primary sensory areas **(Shipp, 2005; Shipp, Adams and Friston, 2013)**. Here, by analyzing laminar recordings in the motor cortex of a monkey performing a complex visuomotor task, we provide novel evidence that challenges the idea of localizing different types of task-related information in distinct specialized layers. Instead, we find that these information types are simultaneously encoded within alternative low-dimensional spaces of coordinated neural activity, all generated by the interactions across all layers. At specific moments in time, however, task-related information becomes dynamically concentrated in either superficial or deep layers, and traveling waves of information emerge against a background of more unstructured, generic fluctuations of activity spanning the entire column.

Motor cortex represents a privileged place for investigating complex multilaminar processing. Far from being a pure *output structure*, this precentral region acts as a functional hub within diverse distributed networks, which provide it with direct access to both sensory and cognitive inputs to guide behavior **(Wise, 1999; Rizzolatti and Luppino, 2001; Battaglia-Mayer and Caminiti, 2019)**. Integrating these heterogeneous signals relies therefore necessarily on the anatomical constraints and functional connections within the cortical column. In contrast to primary sensory areas, motor cortex displays a less strict segregation of afferent and efferent projections, enhanced intra-laminar connectivity and an almost indistinguishable layer IV **(Shipp, 2005; Shipp, Adams and Friston, 2013; García-Cabezas and Barbas, 2014; Godlove *et al*., 2014; Ninomiya *et al*., 2015; Borra *et al*., 2021)**.

To investigate the computational mechanism emerging from this *weakly laminated* vertical structure, researchers recorded electrophysiological signals from the premotor cortex in monkeys during complex tasks. Based on single-unit activity, Chandrasekaran and colleagues **(2017)** proposed a segregation scheme of information processing, with decisions preferentially encoded in superficial layers and movement-related signals in deep layers. Conversely, Opris and colleagues found that neurons were modulated across all layers, with behavioral performance relying on synchrony between neuron pairs at different depths within cortical *minicolumns* **(Opris *et al*., 2011, 2015; Opris, 2013)**. These studies thus propose two different mechanisms of cortical multiplexing: a layer-based versus a column-based scheme of computation.

When task complexity increases, the interpretability of single-unit responses decreases and their heterogeneity increases, as individual neurons often exhibit mixed selectivity and encode simultaneously multiple task variables with different encoding schemes **(Rigotti *et al*., 2013)**. However, interpretability is rescued by considering the joint activity of multiple neurons as describing a trajectory in a space of the system’s dynamic configurations **(Mante *et al*., 2013; Zhang *et al*., 2025)**. Lower-dimensional manifolds -or vector subspaces in a simpler linear framework-parameterized by a few latent variables can be identified **(Churchland *et al*., 2012; Shenoy, Sahani and Churchland, 2013; Vyas *et al*., 2020; Genkin *et al*., 2025)** in which dynamic trajectories unfold and separate meaningfully according to behavioral features **(Gallego *et al*., 2017; Perich, Narain and Gallego, 2025)**, consistently across time and individuals **(Gallego *et al*., 2020; Safaie *et al*., 2023)**. Furthermore, recent findings indicate that neural subspaces exhibit flexibility, being both *reused* to support the same function over time, and *recycled* to encode novel variables **(Driscoll et al., 2024; Tafazoli et al., 2026; Wentz et al., 2025; Yang et al., 2019)**.

In the present work, we examined how the motor cortex exploits its intertwined laminar structure to generate latent representations. The cortical layers define a *laminar space* (Figure 1), where activity forms vectors of instantaneous multilaminar states. By tracking task-relevant patterns within this space, we agnostically assessed whether these informative subspaces are localized in depth -suggesting individual layers act as functional units- or instead span multiple depths, pointing to the full column as the functional unit. Furthermore, we investigated whether these coding spaces were *reused* over time to maintain stable representations of the same variables, or were flexibly *recycled* for representing alternative variables at different task epochs **(Wentz *et al*., 2025)**.

**Figure 1.**
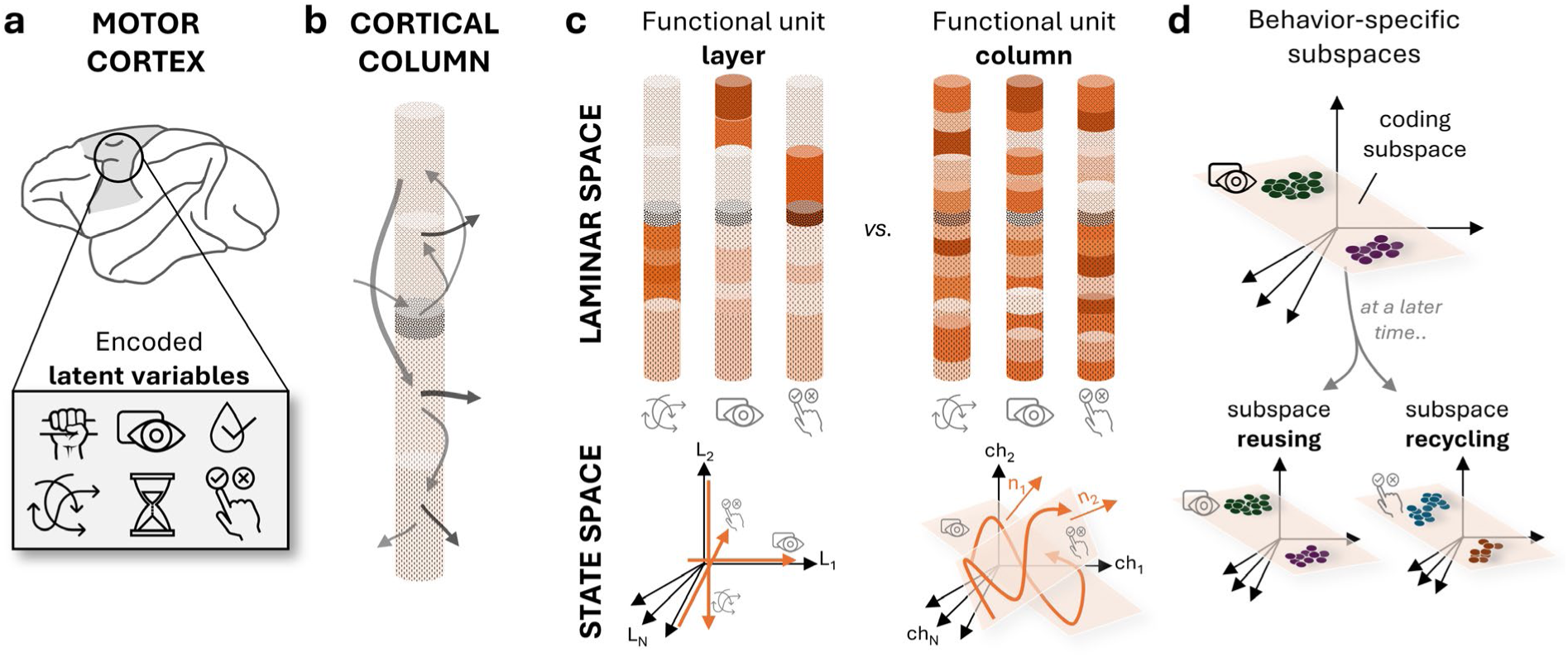
Hypothesis: Laminar functional units in the motor cortex enable multiplexing. **a**, The motor cortex encodes multiple task variables (e.g., force, visual cues, reward, direction, time, choice). **b**, The laminar organization of the cortex provides a substrate for multiplexing. **c**, Two possible multiplexing mechanisms are illustrated. Left: Layer segregated multiplexing: different layers act as independent functional units, each preferentially encoding specific variables. In state-space, coding subspaces would be defined by channels within the same layer, and movement along individual axes would describe the encoded variables. Right: Column distributed multiplexing: the entire column acts as a functional unit, with different combinations of channels contributing to the encoding of different variables. In this case, coding subspaces would be formed by channels spanning the entire column, and trajectories within these low-dimensional spaces would capture variable encoding. **d**, Coding-specific subspaces could be reused whenever the same variable is being encoded or recycled to encode distinct parameters over time.

We found that information multiplexing, manifested as the simultaneous representation of two distinct latent variables by a same circuit, was supported by the coexistence of different low-dimensional subspaces arising from different combinations of layers within the full-dimensional laminar space. The fact that complex multilaminar coordination patterns were reused consistently in a behaviorally related manner suggests that information processing relies more on integration than strict segregation across layers. Yet, we found that the encoding subspaces themselves evolved over time, giving rise to a structured flow of task-related information that smoothly alternated between being predominantly represented in superficial and in deeper layers. Laminar localization of information could therefore emerge at times, but only as transient episodes within a broader dynamic that always unfolded over the entire laminar space and varied across recording sites.

## Results

### Multi-unit activity during delayed match-to-sample task

To study the spatial and temporal structure of laminar coding subspaces in the motor cortex we analyzed laminar population activity across layers during visuomotor behavior in monkeys, in a task requiring simultaneous encoding of cue identity and direction.

Specifically, two macaques were trained to perform a temporally predictive delayed match-to-sample (DMS) task (Figure 2-a). This complex task requested them to transform the information about the identities of the cues (color condition) into the reaching direction to be performed after the GO. They had to hold in memory the SEL cue identity and match it with the appropriate SC to determine the correct movement direction.

**Figure 2.**
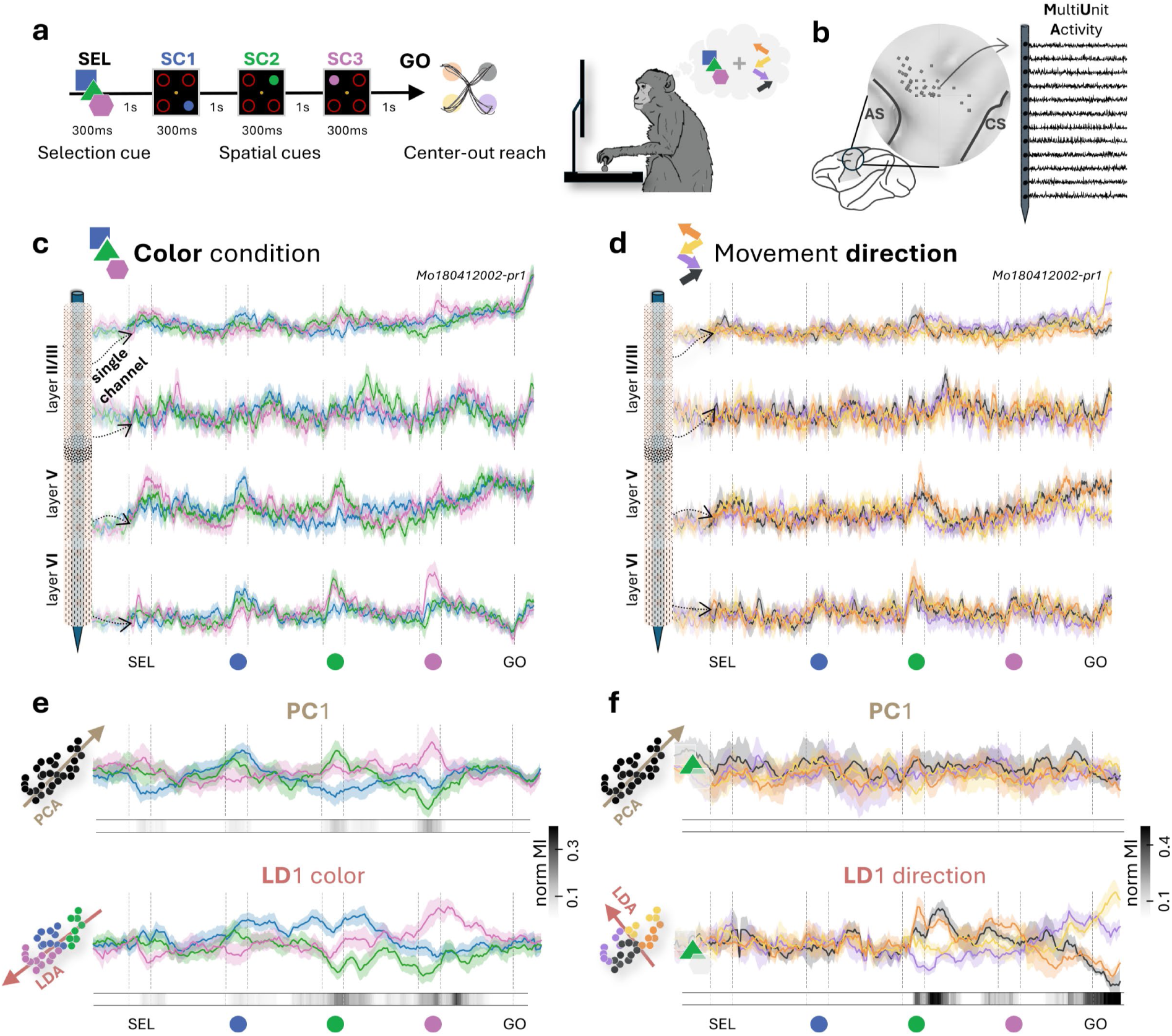
Laminar dynamics in the motor cortex encode task-related variables. **a**, Monkeys performed a temporally predictive delayed match-to-sample reaching task. In each trial, a color-coded selection cue (SEL) indicated the color to attend to (sample). Then, three differently colored spatial cues (SCs) -only one matching the color of SEL- and a GO signal were presented. After GO, the monkeys had to perform a center-out reaching movement to the memorized location of the color-matching SC (match) to obtain a liquid reward. The SCs were always presented in the same order (SC1 blue – SC2 green – SC3 pink), but their location was random (1 out of 4 possible diagonal locations). Each delay lasted 1s and each visual cue 300ms. This task requested monkeys to transform the cue identity (color condition) into a movement direction (match SC). **b**, Left: Laminar sites plotted on the MRI-based cortical surface reconstruction of monkey T (anterior: left, posterior: right); AS: arcuate sulcus, CS: central sulcus. Right: Example multi-unit activity (MUA) recorded with a linear multielectrode array at one site; 12 selected channels from a 19-contact probe, preserving depth organization. **c**, Trial-averaged MUA separated by color-condition (SEL identity) for one channel per layer, spanning the cortical column. Shaded area: 95% confidence intervals (CI). **d**, Same as in (c) for green (green-SEL) trials separated by movement direction; black: upper right, purple: lower right, yellow: lower left, orange: upper left. **e**, Neural activity projected onto the first axis of Principal Component Analysis (PCA, top) and Linear Discriminant Analysis (LDA, bottom), calculated in 200ms windows with 25ms shift. Time-resolved projections were concatenated to reconstruct the trial. Color condition (SEL identity) means are displayed as color-coded lines. Shaded area: 95% CI. Gray bars below indicate significant mutual information (MI), with intensity reflecting magnitude; white indicates non-significant bins. **f**, Same as in (e) for green trials separated by movement direction. (**c-f**), Example site: Mo180412002-pr1.

After monkeys had learnt the task, we recorded the neural activity during task performance using acute multi-channel linear (laminar) arrays in the primary motor/premotor cortex. This gave us access to different cortical columns along the cortical sheet (Figure 2-b). We then measured neuronal population activity across layers computing the Multi-Unit Activity (MUA) signal on each individual channel, reflecting the fluctuations of the local population spiking nearby each contact **(Stark and Abeles, 2007)**.

Trial-averaged neural activity in individual channels across all layers showed modulations with respect to the two main variables encoded during the task: 1) color condition (selection cue identity) and 2) movement direction (Figure 2-c, d). Such modulations were, however, weak, often not significant, and heterogeneous in their exact depth across the different cortical sites.

### Dimensional reduction of population activity dynamics

To filter out irrelevant variability, we performed a dimensional reduction of population activity recorded simultaneously across all channels, by applying two different linear methods: principal component analysis (PCA) and linear discriminant analysis (LDA). While PCA captures the directions of maximum variability, LDA identifies axes that specifically maximize separation across the probed conditions, thereby defining coding subspaces in which trial trajectories for those conditions (SEL color or target direction) are better separated. These models were estimated in a time-resolved manner along the trial, and neural activity was projected at every time point onto the current first axis in each case.

When separating trials by color condition, the single-trial PCA projections showed time courses resembling those observed at the single-channel level. Differences between color conditions -quantified by significant mutual information (MI)-arose mainly around the presentation of visual cues (Figure 2-e, top). In contrast, LDA projections were able to track color information throughout the trial, reflected in longer-lasting periods of significant MI, extending into the delay periods (Figure 2-e, bottom).

For movement direction, PCA projections did not capture direction-specific dynamics, with not significant MI at any point (Figure 2-f, top). LDA projections, however, still diverged according to target direction, starting from the moment in which the valid spatial cue (color-matching SC) was presented, i.e. when information about the target movement direction first became available (see example session for green trials, in which valid direction was presented at SC2, Figure 2-f, bottom).

Therefore, the directions of maximum variability do not align with the coding axes for task-related variables. This indicates that most inter-trial variability is task-unrelated -or at least not directly linked to color and target direction encoding- and that supervised approaches such as LDA are required to better disentangle the spatiotemporal organization of task-relevant activity fluctuations in laminar space.

### Revisiting coding subspaces along the trial

Since LDA latent dimensions effectively captured the dynamics of the two main task-related variables, we next investigated the contributions of different layers to generate these coding subspaces and how stable they were throughout the trial.

At every time, an LD direction is recomputed and defined by a vector of entries over every channel depth in laminar space, providing the weighted and signed contribution of each channel’s activity to the considered coding subspace. To visualize channel contributions, we plotted these dynamically-changing weights for LD1_color_ and LD1_direction_ (Figure 3-a, color: top, direction: bottom; PC1 weights are shown for comparison in Suppl.Figure 1-a) in a matrix form where each vertical slice represents the LD at a different time. The primary coding axes for both color and direction displayed strong heterogeneity in channel contributions within and across layers, alongside remarkable temporal changes.

**Figure 3.**
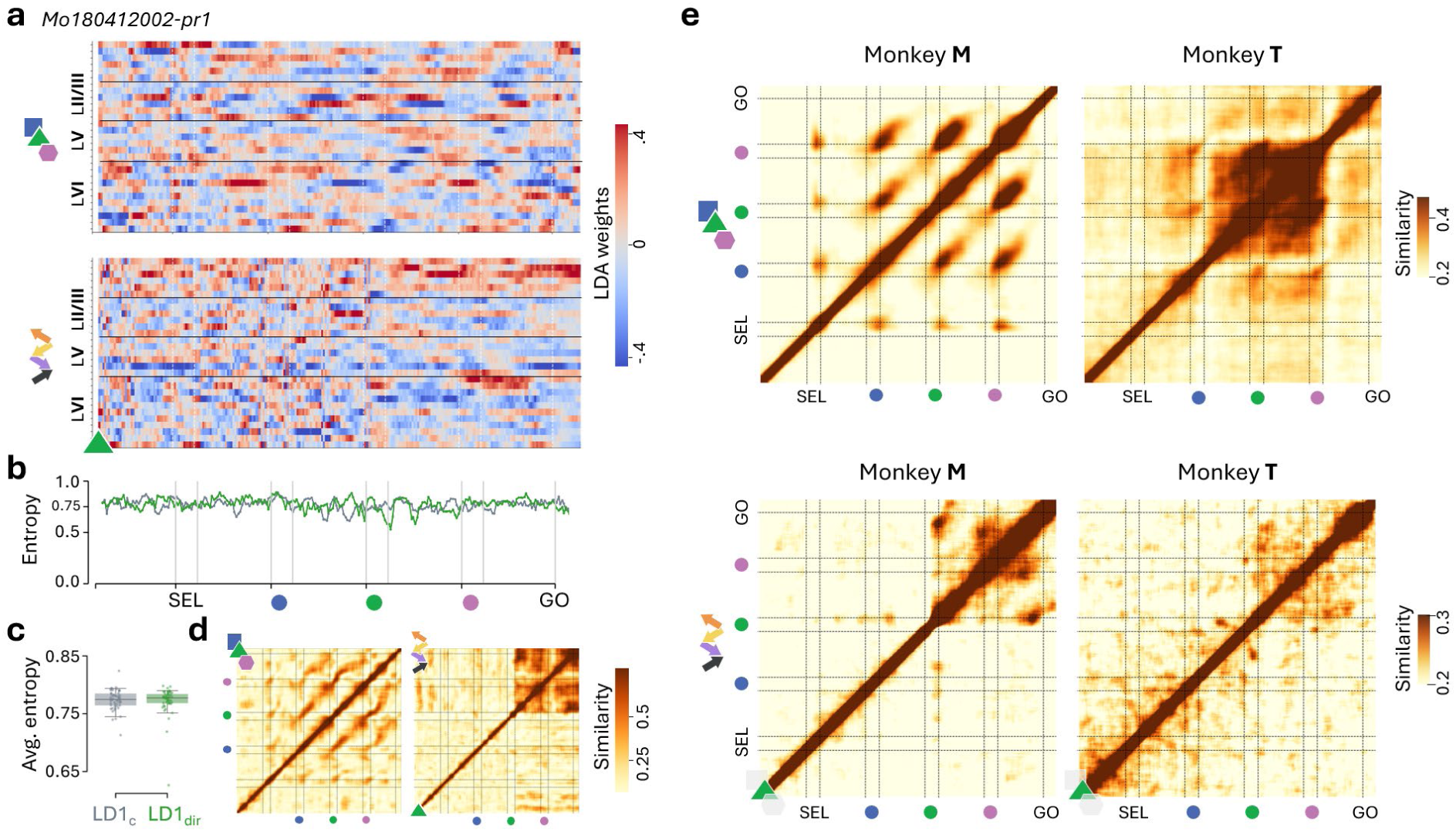
Laminar coding spaces are revisited along the trial. a, Heatmaps displaying the weights of all time-resolved LD1_color_ (top) and LD1_direction_ (green trials, bottom) models for the example site. Horizontal lines indicate the boundaries of cortical layers; x-axis: trial time, y-axis: channels. b, Time-resolved Shannon entropy of weight vectors in (a). Values of zero indicate all weight in a single channel and one equal contribution from all channels. x-axis: trial time. c, Distribution of the average entropies of LD1_color_ (gray) and LD1_direction_ (green) weights across all recorded sites; whiskers indicate 5^th^-95^th^ CI. d, Cross-temporal map of absolute cosine similarity across pairs of weights in (a); left: LD1_color_, right: LD1_direction_; x-axis, y-axis: trial time. Maps represent the average across 50 similarity maps built on resamples of training data. e, Same as in (d) but averaged per monkey across all sites; left: Monkey M (n=27 sites), right: Monkey T (n=14 sites), top: LD1_color_, bottom: LD1_direction_ in green trials. (a-d), Example site: Mo180412001-pr1.

Visual inspection of the computed LDs did not reveal any marked concentration of relevant weights at specific depths. To quantify the degree of laminar spread in the weights defining the coding subspaces, we used a Shannon entropy metric such that a value of one indicates equal contributions from all channels, while a value of zero indicates that the LD is perfectly aligned with activity in just a single channel. Here, all subspaces (both variables at all times) exhibited an entropy level around 0.75, implying that information was highly distributed along the column (Figure 3-b); besides the shown example session, similarly large values were generally observed across all sites recorded (Figure 3-c).

We then checked whether these spatially distributed laminar coding subspaces were retrieved at different times. To this end, we performed a recurrence analysis to quantify the similarity between LDs extracted at any possible pair of time points along the trial (in terms of a bounded cosine similarity, where 0 means orthogonality of vectors and 1 perfect alignment). Clear task-related cross-temporal patterns emerged for both color (Figure 3-d, left) and direction (Figure 3-d, right), shown by the recurrence matrices for the example session displayed in Figure 3-(a, b), and especially from the recurrence matrices in Figure 3-e, averaged over all recording sites for each of the monkeys. Blocks and stripes in the recurrence matrices denote phenomena such as transient stabilization of a coding subspace (a block along the diagonal; larger blocks indicate longer stabilization) or repeated recruitment of previously used coding subspaces (off-diagonal stripes or blocks, denoting enhanced inter-time similarity of LDs).

Focusing first on the color-condition encoding, and specifically on the dynamics LD1_color_ axis, we observed in the session-specific recurrence matrix (Figure 3-d, left) the emergence of off-diagonal stripes and blocks, reappearing when new spatial cues were presented. These persisted even after averaging across all recording sites, being particularly evident for the first monkey (Figure 3-e, top left) and more blurred for the second (Figure 3-e, top right).

The off-diagonal blocks observed in Figure 3-e (top) indicate that the coding subspaces carrying information about color conditions in the epochs following SC2, and SC3 largely overlap with the coding subspace already recruited at SC1 (and, for monkey M, even with that engaged earlier at SEL cue). At the single-session level (Figure 3-d, left), the off-diagonal stripes connecting these block locations reveal an even tighter temporal replication of the precise sequence of color-condition coding subspaces, recruited throughout the delay periods between two consecutive cue presentations.

Considering next the encoding of target direction, the dynamics of the direction-related LD1 differed from those of the color subspaces. The target direction cannot be determined until the spatial cue matching the selection cue color -hereafter the matching cue-is presented (e.g., SC2 if the color pre-cued at SEL is green). Figures 3-d (right, single representative session) and 3-e (bottom, averaged across sessions) show recurrence matrices for the LD1_direction_ in green trials, while analogous matrices for blue and pink trials are provided in Suppl.Figure 1-b. Before the presentation of the matching cue, the direction-coding subspaces evolved in a largely random manner. After the matching cue, however, the direction-coding space stabilized, as indicated by a darker diagonal block appearing immediately after the cue (compare Figure 3-e, bottom, for green trials where the matching cue is SC2, with Suppl.Figure 1-c for blue and pink trials, where the matching cues are SC1 and SC3, respectively). Notably, we observed a spatial alignment between the direction-coding subspace during the delay following the matching cue and that preceding the GO signal, consistent with the notion of a preparatory subspace that is revisited prior to movement initiation.

While the averaged recurrence LD1_direction_ matrices for both monkeys preserved several salient elements of single-session matrices, some differences were observed across the two animals. These differences likely reflect previously reported differences in spontaneous micromovements **(Nougaret *et al*., 2024)**, more transient in monkey M than in monkey T (see discussion). Specifically, in Monkey T, the stability of the direction-coding subspace was weaker but persisted longer after the matching cue, extending up to movement onset following GO. By contrast, in Monkey M, stability was more intermittent, but the pre-GO and post-match delay coding subspaces showed stronger alignment. Irrespective of subtle differences between monkeys and encoded variables, these results indicate that the dynamics of coding subspaces were strongly constrained by the task structure.

### The functional subspace encoding cue identity is recycled as a match subspace

Given that similar color-encoding subspaces were revisited during visual cue presentations (Figure 3-e, top), we first analyzed how color-related information was specifically represented within these subspaces by projecting single-trial laminar MUA onto one of them. As a reference, we chose the subspace encoding cue identity during SEL. To perform the task, monkeys had to select the spatial cue matching the selection cue (SEL) color. Therefore, we hypothesized that the laminar space encoding the cue identity (color) during SEL could either be reused during spatial cues to again encode the same cue identity/color (H_1_) or recycled as a match space (H_2_) (Figure 4-a). These two hypotheses yield specific predictions about how the cue-identity space at SEL should generalize:

**Figure 4.**
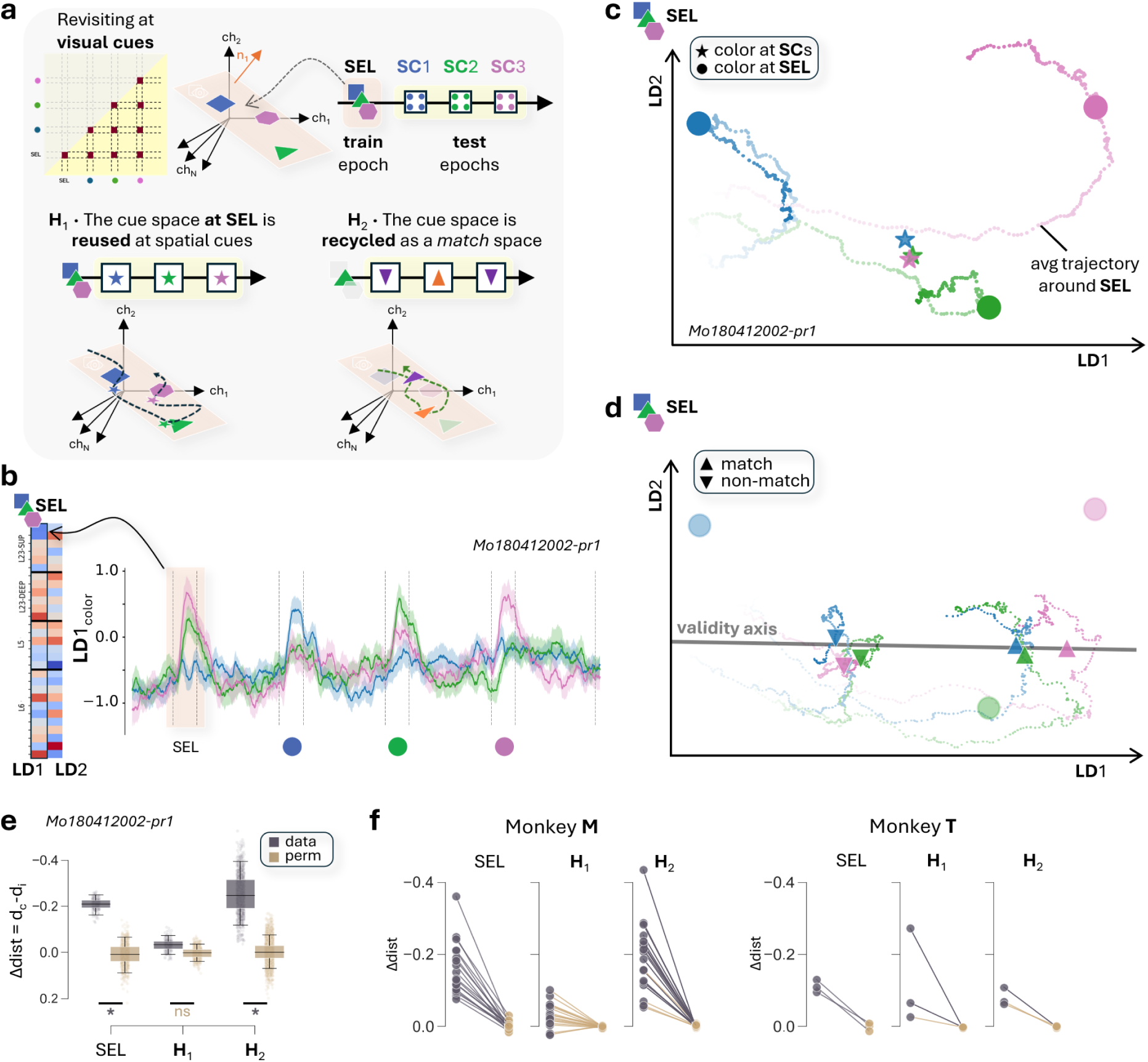
The coding space for cue identity is functionally recycled as a match space. a, Schematic illustration of two hypothesis: cue identity subspace reuse for color encoding (H_1_) or recycling for match encoding (H_2_). Cue identity space revisiting was tested based on cosine similarity between SEL and SC epochs (upper left). An LDA model was trained at the selection cue (SEL, train epoch) to separate color conditions (blue, green and pink). Neural activity during spatial cues (SCs, test epochs) was then projected into this model. Bottom left: H_1_ (color encoding). Stars represent trial-averaged states for each SC, color-coded by SC color (SC1: blue, SC2: green, SC3: pink). Bottom right: H_2_ (match encoding). Upwards orange triangles indicate trial-averaged states for matching trials (e.g. green trials at SC2) and downwards purple triangles states for non-matching trials (e.g. green trials at SC1/SC3). b, Left: Channel weights for LD1/LD2 at SEL. Right: Trial-averaged projections on LD1 over time, separated by SEL color; line: mean, shaded: 95% CI, x-axis: trial time. c, Condition-averaged trajectories around SEL in the LD1-LD2 space, color-coded by SEL color; decreasing transparency indicates elapsed time, from 50ms after cue onset until cue offset. Large colored dots represent the state of maximum cross-condition variance during the train epoch. Color-coded stars describe trial-averaged projections on SC1 (blue), SC2 (green) and SC3 (pink). d, Same LD1-LD2 space as in (c); with trial-averaged projections color-coded by the SC color; decreasing transparency indicates elapsed time, from 50ms after cue onset until cue offset. Upwards triangles: matching trials, downwards triangles: non-matching trials; (e.g. match: green upwards triangle for green trials at SC2; non-match: green downwards triangle for blue and pink trials at SC2). Gray line: validity axis. e, Distribution of the single-trial distance metric (Δ_dist_), computed as the difference between distance to correct (d_c_) and incorrect (d_i_) clusters; dark gray: data, light brown: permutations. Analysis was repeated across 200 train/test splits and 1000 permutations to test for SEL space (train epoch), H_1_ (color encoding) and H_2_ (match encoding). f, Summary across sites. Left: Monkey M, right: Monkey T. Only sites with a significant SEL space (Δ_dist_ data < 95^th^ percentile of permutations) were included. Dots: mean Δ_dist_ for each test: SEL, H_1_, H_2_ (both data and permutations); dark connecting lines: significant results after Bonferroni correction. (b-e), Example site: Mo180412002-pr1.

Under H_1_, the cue-identity subspace would be functionally reused as a *pure color-coding space*, such that the population state represents the color displayed on the screen, regardless of whether the currently shown cue is a distractor or the matching spatial cue. In this case, the codes defined for blue, green, and pink cues at SEL would be reused for all trials at SC1, SC2, and SC3, respectively. Consequently, trajectory snippets for single trials associated with different colors, when projected onto the reference coding subspace, should converge toward distinct split locations characteristic of each color, independently of when that cue was presented (Figure 4-a, bottom left).

In contrast, H_2_ proposes that the cue-identity subspace at SEL is functionally recycled as a *match space* during the presentation of the SCs (Figure 4-a, bottom left). Under this scenario, what is encoded in the reference subspace at later SCs is not the color displayed on the screen, but instead whether the cue matches the color originally presented at SEL. In other words, the code no longer represents cue color per se, but rather its validity -i.e., its pertinence for extracting directional information. Accordingly, the projected multilaminar neural activity snippets would converge toward just two split locations, corresponding to “valid” or “non-valid,” irrespective of the specific color shown (e.g., blue-trial snippets at SC1, green-trial snippets at SC2, and pink-trial snippets at SC3 would all converge, after projection onto the reference subspace, toward the common spot encoding the cue as “valid”).

To test these hypotheses, we first built the reference coding subspace for cue identity at SEL. Because three colors had to be discriminated, such a subspace (LD_color_) is spanned by two discriminant vectors whose weights over different columnar depths -the different channels in the laminar probe-are shown to the left of Figure 4-b for a representative site. On the right of Figure 4-b, we then show the average projections of trials for different color conditions (SEL color) over the first axis of this reference space. We observed that the condition-averaged activity presented a peak corresponding to each visual cue. Importantly, the amplitudes of the peaks at SCs were higher whenever the cue matched the color condition, providing initial evidence that cue color is processed differently across trial epochs. In a purely color-coding scenario, the amplitudes of the three peaks at SCs should scale with their respective color amplitudes during SEL, thus preserving the same color code.

An even stronger indication against the pure color space hypothesis (H_1_) comes from bidimensional projection on the plane spanned by the first and second LD axes (Figure 4-c). The big dots correspond to the moment of maximum cross-condition variance in the train epoch, colored based on the SEL cue identity; the small colored dots represent the condition-averaged projected trajectory snippets during SEL. We then projected in this same reference space trajectory snippets during a small window surrounding SC1, SC2 and SC3 presentations and marked as colored stars (respectively blue, green and pink) the average barycenter of these projected test snippets. We observed that these stars clustered in the middle of the plane, far from the split locations coding SEL cue identity. Therefore, we conclude that this reference subspace was not reused to encode color, since colored stars did not overlap with colored dots.

To test for H2, we again projected laminar MUA on the same reference subspace, but according to a different, validity-based split. Specifically, in Figure 4-d we show averaged trajectory projections and average barycenters for trials at SC1, SC2 or SC3, color-coded by spatial cue color and further separated by validity condition. Upward triangles correspond to barycenters of MUA trajectory snippet projections of matching cues (e.g. around SC1 for blue trials). Downward triangles correspond to the barycenters in non-matching condition (e.g. around SC1 for green and pink trials). For reference, we also show the colored-dots corresponding to the original color separation at SEL (same as Figure 4-c). Here, we can observe a clear distinction between two clusters, with barycenters grouped according to validity (match vs non-match conditions) of the currently shown cue, irrespective of its color, precisely in line with the predictions of H2 and in contrast with the ones of H1. We could thus find a one-dimensional axis embedded in the originally bidimensional reference coding plane which was able to separate valid from invalid when the data was projected onto it. This corresponds to a case of recycling the original coding subspace for cue identity, i.e. a transformation of representation beyond identical reusing.

To validate these observations in single trials across cortical sites, we implemented a quantitative statistical analysis. Specifically, for each recording site we constructed a site-specific reference color coding subspace at SEL and determined, within this space, the barycenters of the categories to be discriminated (blue vs. green vs. pink at SEL; valid vs. invalid at later SCs). Then, in each trial, we computed the distance of projected snippets to these category barycenters. The rationale is that a decoder generalizes appropriately if a trial’s projection lies significantly closer to the correct category barycenter than to incorrect one(s). Statistical significance was assessed via permutation testing. We quantified this effect using the difference Δdist between distances to incorrect versus correct categories under different conditions.

Figure 4-e illustrates statistical testing for the same site as b-d. As expected, projected snippets at SEL clustered significantly closer to the correct color-category barycenters than to the others, with cross-validated Δdist consistently above chance, supporting the existence of a cue-identity encoding space at SEL (Figure 4-e, left). When projecting snippets from later SCs, the color-based separation was no longer significant (i.e., H1 fails), as Δdist did not exceed chance levels (Figure 4-e, center). However, when discriminating between the two categories “valid” vs. “invalid”, Δdist for projected snippets from later SCs rose again consistently above chance (i.e., H2 was supported for this recording site; Figure 4-e, right). Results across sites and monkeys are summarized in Figure 4-f.

The recycling of the color-identity subspace at SEL differed between the two monkeys. In monkey M, where the SEL color subspace revisiting at later SCs was clear already from the site-averaged similarity map (Figure 3-e, top left), 18 out of 27 sites presented a significant cue identity space at SEL. From those, H1 was rejected for all of them (light lines, not significant) and H2 was confirmed in 14 out of 18 sites (dark lines, significant) (Figure 4-f, left). This indicates that, for this monkey, the cue identity space at the selection cue was *recycled* as a match-to-sample space at the spatial cues, enabling the transformation of the cue information into a match/non-match representation.

In Monkey T, by contrast, we already noted that the average recurrence matrix for color-coding LDs (Figure 3-e, top right) lacked the off-diagonal block indicating cross-similarity between SEL and spatial cues. This is explained by the fact that, for most sites in monkey T, we were unable to identify a color-identity coding subspace at SEL. Indeed, such a subspace significantly discriminating cue color identity at SEL was found in only 3 out of 14 sites. In the rare cases where these cue-color subspace existed, 2 of the 3 continued to operate as color-coding spaces (consistent with H1), and 1 of them also simultaneously acted as a validity-coding space. In conclusion, for monkey T, analysis of the recycling of cue-identity coding spaces at later SCs is largely ill-posed, as the reference subspace itself generally fails to exist. Instead, the data suggests the presence of a stable representation of the color condition across all task periods (Suppl.Figure 2), consistent with other behavioral differences observed in this animal (see discussion).

### The match space is functionally reused at the spatial cues

Although Monkey T did not exhibit a clear cue-identity space at SEL that could be later recycled, the recurrence matrix (Figure 3-e, top right) nonetheless indicated the presence of persistent color-coding subspaces. We therefore examined whether an alternative reference coding subspace, extracted at a different task moment, might instead be functionally reused. Specifically, we focused on color coding subspaces at a later cue (e.g. SC1) and hypothesized (H_1_) that it could be reused at other cues (e.g. SC2 and SC3) to encode cue validity (Figure 5-a).

**Figure 5.**
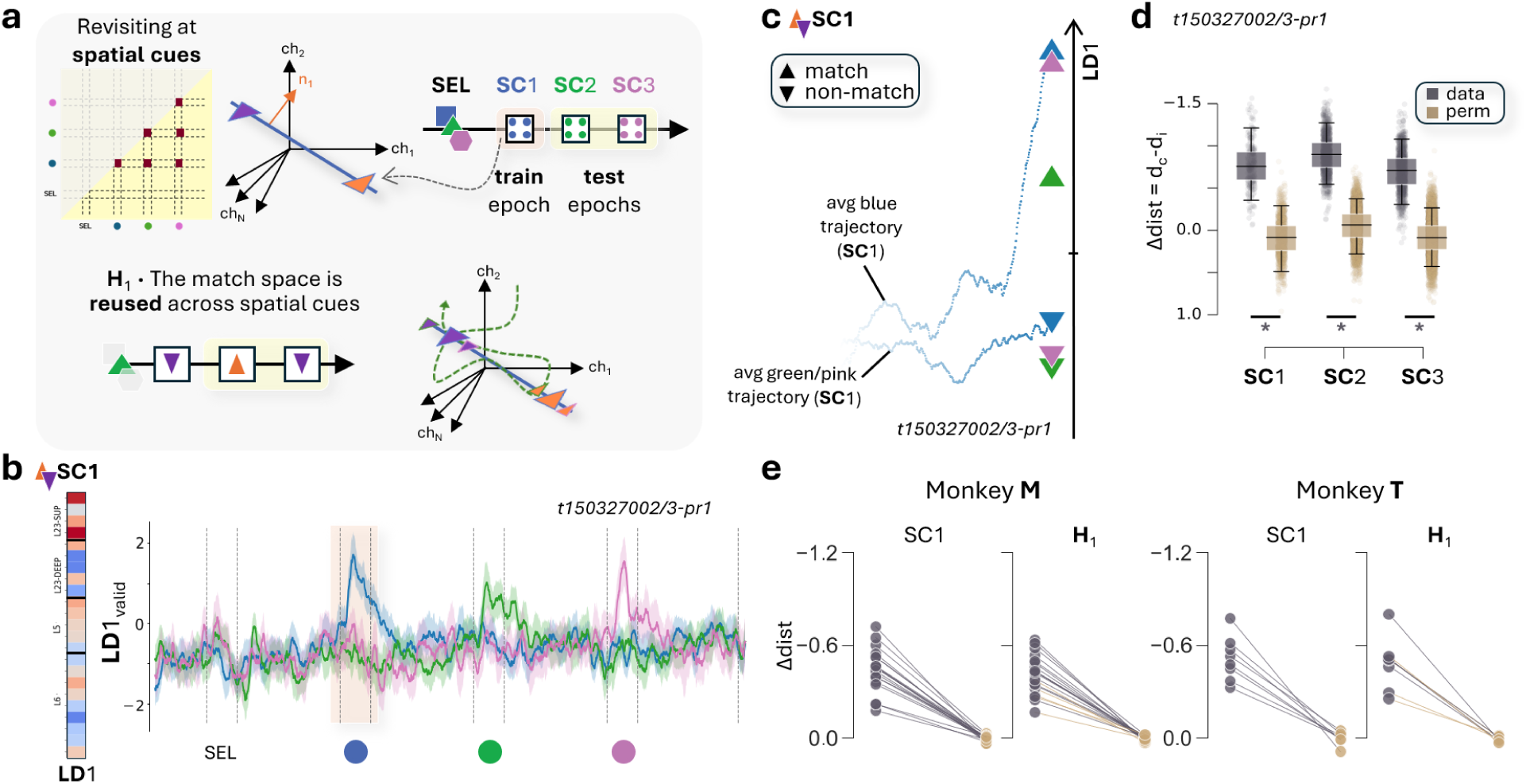
The match space is functionally reused across spatial cues. a, Schematic illustration of the match space reusing hypothesis across spatial cues (H_1_). Match space revisiting was tested based on cosine similarity among SCs (upper left). An LDA model was trained at SC1 (train epoch), to separate match from non-match (blue vs. others). Neural activity during the other spatial cues (SC2/3, test epochs) was projected into this model. Bottom: H_1_ (match encoding). Upwards triangles indicate trial-averaged states for matching trials (e.g. green at SC2) and downwards triangles states for non-matching trials (e.g. green at SC1/SC3). b, Left: Channel weights for LD1 at SC1. Right: Trial-averaged projections on LD1, separated by SEL color; line: mean, shaded: 95% CI, x-axis: trial time. c, Condition averaged trajectories around SC1 in the LD1 axis; decreasing transparency indicates elapsed time, from 50ms before cue onset until the time of maximum cross-condition variance (around SC1). Blue triangles represent the maximum cross-condition variance (match/non-match) during the train epoch. Color-coded triangles describe the trial-averaged projections on the other SCs color coded by SC color. Upwards triangles: matching trials, downwards triangles: non-matching trials. d, Distribution of the single-trial distance metric (Δ_dist_), computed as the difference between distance to correct (d_c_) and incorrect (d_i_) clusters; dark gray: data, light brown: permutations. Analysis was repeated across 200 train/test splits and 1000 permutations to test for SC1 match space (train epoch), SC2/SC3 match spaces (test epochs); stars denote significant difference. e, Summary across sites. Left: Monkey M, right: Monkey T. Only sites with a significant SC1 match space (Δ_dist_ data < 95^th^ percentile of permutations) were included. Dots: mean Δ_dist_ for each test: SC1, H_1_(SC2/SC3) (both data and permutations); dark connecting lines: significant results after Bonferroni correction. (b-d), Example site: t150327002/3-pr1.

To test this hypothesis, we constructed a new reference LD axis (e.g., at SC1) to discriminate between valid and invalid trials. Since the categories to be separated were now only two (valid vs. invalid), the extracted coding subspaces were explicitly one-dimensional. As before, the weights of the discriminant vectors were not concentrated in a few layers but were broadly distributed across cortical depths (see example site for Monkey T in Figure 5-b, left).

When examining the time courses of trial projections onto this axis, averaged by color condition, we observed a marked peak at the matching cue, already suggesting subspace reusing (Figure 5-b, right). An alternative representation is shown in Figure 5-c, where we illustrate the average time evolution of trajectory snippets around SC1, projected onto the reference LD axis for blue (valid) and green/pink (invalid) trials. These projections converged to distinct barycenters (upward vs. downward blue triangles), well separated along the reference LD. Similarly, trajectory snippets around the other SCs converged to distinct barycenters (upward vs downward triangles) depending on their validity. Across all cases, barycenters clustered along the reference LD axis according to validity, irrespective of color identity or task epoch.

We next quantified discrimination at the single-trial level using the Δdist metric. Individual trials were separated well above chance not only around SC1, where the reference space was constructed, but also when reusing the same space to discriminate validity at later cues SC2 and SC3 (Figure 5-d, example session). This ability generalized well across monkeys and recording sites (15/20 in M; 4/8 in T; Figure 5-e).

In summary, both monkeys displayed a visual subspace encoding cue validity - independently of cue identity- that was functionally reused during spatial cue presentations. In Monkey M, this subspace was recycled from the cue-identity space, whereas in monkey T it was restricted to spatial cues. In both cases, this subspace might serve as a gating mechanism, determining whether incoming spatial information is retained or ignored.

### Two distinct subspaces -visual and motor-encode target direction

After the encoding of cue identity and validity, we investigated the encoding of direction. Specifically, we hypothesized that a coding subspace for the target direction existed and was reused across trials whenever the correct SC was presented (H_1_). Note that target direction refers to the movement direction to be performed at GO, which can only be inferred at the time of the matching spatial cue presentation. This distinction will become important when considering the encoding of distractor directions, also visually shown, but not task relevant.

If such a coding subspace existed (H_1_), the same code for target direction would be preserved across spatial cues (Figure 6-a). The directional code would be maintained when building a subspace in one cue and projecting the activity around the others (e.g. building the subspace on green trials at SC2 and projecting blue trials at SC1 or pink at SC3, Figure 6-a, bottom).

**Figure 6.**
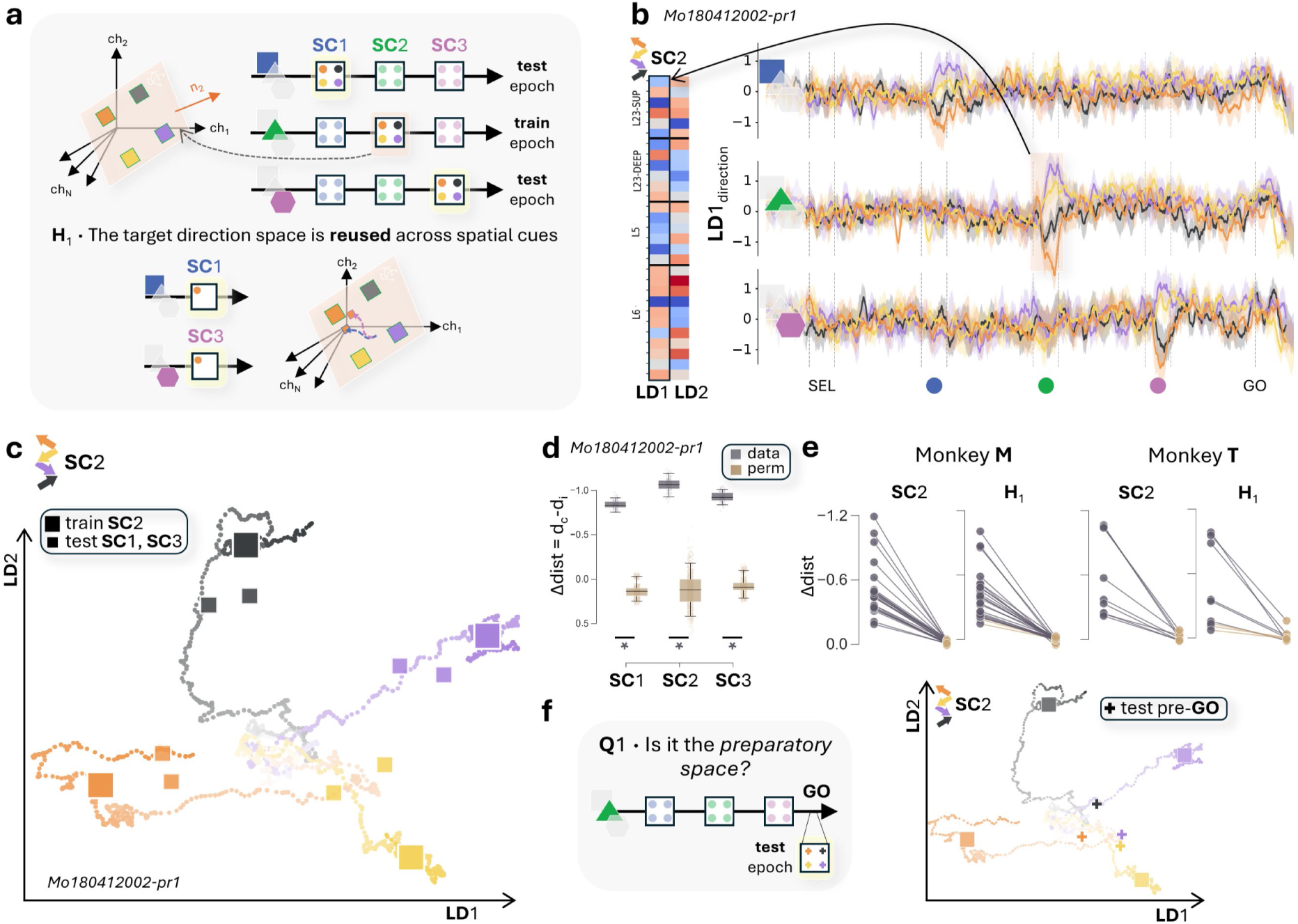
There is a visual, non motor, space functionally reused to encode target direction. a, Schematic illustration of the target direction space reusing hypothesis across spatial cues (H_1_). An LDA model was trained in green trials at SC2 (train epoch), to separate target direction (4 possible). Neural activity during the other spatial cues (SC1/3, test epochs) was projected into this model. Bottom: H_1_ (target direction encoding). Big squares represent green-trial-averaged states for the four target directions at SC2. Small squares depict upper-left direction trial-averaged states for blue condition during SC1 (top) and pink condition during SC3 (bottom). b, Left: Channel weights for LD1/LD2 at SC2. Right: Trial-averaged projections on LD1, separated by movement direction. From top to bottom: blue, green, pink projected trials. Line color: black: upper right, purple: lower right, yellow: lower left, orange: upper left. Line: mean, shaded: 95% CI, x-axis: trial time. c, Condition-averaged trajectories around SC2 in the LD1-LD2 space, green trials color-coded by target direction; decreasing transparency indicates elapsed time, from 50ms before cue onset until cue offset. Large colored squares represent the state of maximum cross-condition variance during the train epoch. Color-coded small squares describe trial-averaged projections per movement direction during test epochs: blue trials projected at SC1 and pink trials at SC3. d, Distribution of the single-trial distance metric (Δ_dist_), computed as the difference between distance to correct (d_c_) and incorrect (d_i_) clusters; dark gray: data, light brown: permutations. Analysis was repeated across 200 train/test splits and 1000 permutations to test for SC2 target direction space (train epoch), SC1/SC3 target direction spaces (test epochs); stars denote significant differences. e, Summary across sites. Left: Monkey M, right: Monkey T. Only sites with a significant SC2 target direction space (Δ_dist_ data < 95^th^ percentile of permutations) were included. Dots: mean Δ_dist_ for each test: SEL, H_1_ (both data and permutations); dark gray connecting lines: significant results after Bonferroni correction. f, Left: Schematic illustration of the target direction space recycled as the preparatory space (pre-GO, test epoch). Right: Same LD1-LD2 space as in (c); with green trial-averaged projections around the GO signal, color-coded by the target direction. (b-d, f), Example site: Mo180412002-pr1.

For illustration, we first built the LD direction subspace for green trials at SC2. The first two discriminant axes of this subspace are shown for an example site in Figure 6-b (left), together with trial projections onto the first discriminant axis (right). Because target direction is inferred at different cues depending on trial color identity, we plotted separate direction-averaged time series for blue, green, and pink trials. As with all other coding subspaces examined so far (and consistent with the entropy analyses in Figure 3-c), the first two discriminants for target direction exhibited weights broadly distributed across cortical depths. The projected MUA time series likewise showed a clear separation between trials with different visually displayed target directions at the matching cue epoch (SC1 for blue, SC2 for green, SC3 for pink), but not at other cues. This observation already is compatible with hypothesis H_1_: direction-coding subspaces constructed at a matching cue would encode target directions at other matching cues, but not generic directions at arbitrary cues (see later section).

The 2D projections onto the plane defined by the two LD axes provided an even clearer visualization of whether a direction-specific code was reused across matching SCs (Figure 6-c). As in previous figures, large squares represent the barycenters of green trials at SC2 (training epoch), with colors indicating the target direction. Direction-averaged trajectories are also shown, highlighting their divergence toward distinct barycenters corresponding to visually presented directions. In addition, we projected trajectory snippets from blue trials at SC1 and pink trials at SC3 onto the same direction-coding reference space built at SC2. Their barycenters are shown as smaller squares using the same directional color code, consistent with H_1_ predictions. Further generalization to other cues is addressed in Suppl.Figure 3-a.

Visual impressions from this example projection plot are once again confirmed by the significance of Δdist analyses at the single-trial level and over distinct sites (see Figure 6-d for example site and Figure 6-e for all sites in both monkeys). In Monkey M, 24 out of 28 sites showed significant generalization at SC2 and 22/24 sites confirmed H1 (Figure 6-e, Monkey M, left). Similarly, in Monkey T, the visual target direction space for SC2 was found in 9 out of 14 sites and 6 out of these 9 confirmed H1 (Figure 6-e, Monkey M, right). Similar generalization performance was observed when constructing the reference space at other matching SCs (Suppl.Figure 3-b). These results demonstrate the existence of a target-direction encoding space that is reused whenever the matching spatial cue is encoded.

When considering direction encoding, a potential ambiguity arises between visual and motor interpretations. At SCs, directional information reflects the screen location of the visual cue, a *visual* notion of direction. Around the GO signal, however, movement is executed (and prepared) in a specific direction, corresponding to a *motor* notion of direction. Which type of direction is represented in the reused subspaces identified above?

To address this question, we projected MUA trajectory snippets recorded just before the GO signal into the reference space constructed at SC2 in green trials, separating trials again by the target direction. The barycenters of these projections are shown as crosses, color-coded as in Figure 6-f (example site). In this session, the crosses did not align with the square clusters, indicating that the code expressed at SCs for visually presented target directions was not recycled in the pre-GO window, where direction reflects motor preparation. The direction-encoding subspaces constructed at SCs, and reused across them, capture specifically *visual* - but not motor-target direction. The single-trial quantification across sites and monkeys confirmed and validated this finding: the visual target direction space at SC2 was not recycled as a preparatory space before the GO (recycling occurring in only 1 out 24 sites in Monkey M and none out 9 sites in Monkey T, Suppl.Figure 5-a, left).

This negative result could have been trivially explained by the absence of a preparatory space, i.e., by a lack of detectable direction encoding prior to the movement. To rule this out, we built a new reference direction-coding subspace in the pre-GO period, constructing cross-validated LDs for the planned movement direction. The majority of sites exhibited a significant preparatory space when trained and tested within this epoch (21/24 in monkey M; 7/9 in monkey T; Suppl.Figure 5-a, right). Thus, two distinct direction-encoding subspaces exist: one for visual target direction, reused across SCs, and another for motor target direction, specifically recruited during the preparatory pre-GO window and not recycled from earlier visual cue epochs.

Finally, we tested whether the visual target-direction encoding spaces also represented the directions of distractor cues presented at non-matching SCs, i.e., visually displayed directions not informative about the final motor output. As shown in Suppl.Figure 4 (a: example site; b: generalization), in Monkey M the directions of distractor cues presented *before* the matching cue (e.g., green distractors in pink trials) were also encoded. However, the directions of distractors *after* the matching cue (e.g., green distractors in blue trials) were no longer encoded. In monkey T, we found no evidence of distractor-direction encoding.

Thus, although the visual direction-encoding subspaces constructed at SCs differ from the one recruited during motor preparation in the pre-GO window, they nevertheless prioritize motor-relevant directional information: visual directions unrelated to the target motor direction are either ignored or discarded once the target information needed for motor preparation has been acquired.

### Coding subspaces in motor cortex have a fine and dynamic laminar organization

After analyzing the reuse and recycling of coding subspaces for both color and direction, we return to our initial question: are the complex operations required to process diverse color and direction information performed by specialized, layer-localized populations, or do they rely on the coordinated activity across multiple layers?

By identifying multiple coding subspaces at successive task moments, we could instantaneously “localize” information in latent space: at a given time, information about a particular feature could be extracted from fluctuations along specific latent directions. However, localization in latent space does not imply laminar localization. The discriminant directions defining these coding subspaces did not show strong weight concentration at a specific depths, instead, weights were broadly distributed across the cortical column. Accordingly, fluctuations along a discriminant axis reflected simultaneous MUA variations across multiple depths, indicating that task-related information necessarily emerged from coordinated multilayer activity patterns. At the same time, subtle heterogeneities in the relative contributions of different depths were sufficient to distinguish these subspaces from one another, thereby opening the possibility of multiplexed coding within the same column. Taken together, our findings suggest that the cortical column, rather than individual layers, operates as the functional unit.

We began by analyzing the similarity of laminar weights across different discriminant directions. First, we tested whether subspaces encoding the same task-related variables were more aligned in the high-dimensional space of laminar vectors. To this end, we measured the principal angle between coding subspaces, with smaller angles indicating greater overlap and similarity between their fine laminar organization **(Knyazev and Argentati, 2002)**.

In both monkeys, the subspaces coding for the same latent variable were better aligned (Figure 7-a), consistent with the reuse findings described above for both cue validity and direction. On the contrary, subspaces computed at the same time-window but coding for different variables presented a significantly reduced overlap (Figure 7-b; 17/27 in Monkey M and 5/14 in Monkey T). These representations, although all broadly distributed across cortical depths and layers, therefore remain distinguishable owing to subtle variations in their non-localized laminar profiles, enabling multiplexed representations.

**Figure 7.**
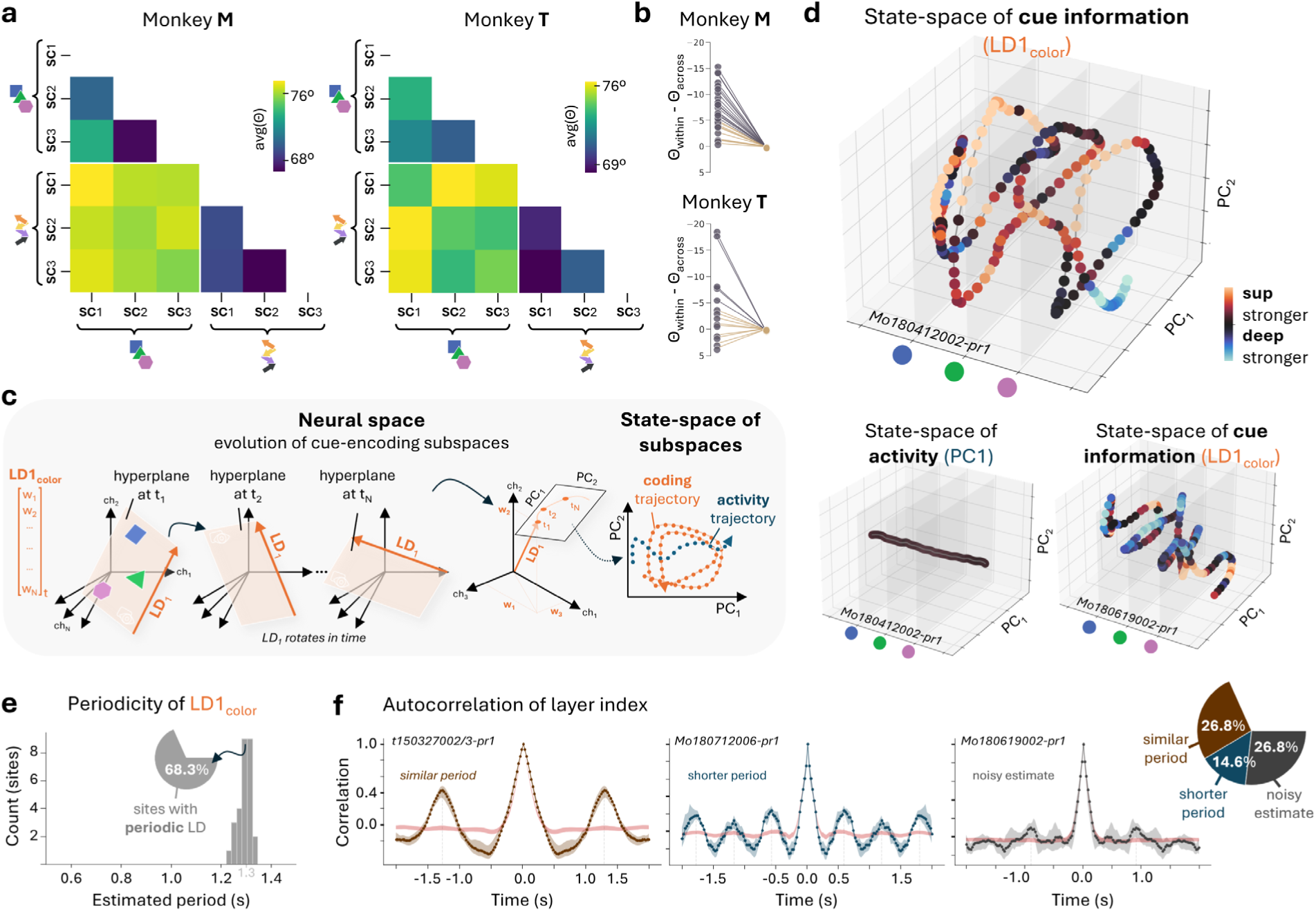
Laminar trajectories of coding subspaces are temporally organized. a, Average subspace angle between pairs of coding hyperplanes across all sites, with darker color for smaller angle. Left: Monkey M, right: Monkey T. b, Statistical summary across sites. Top: Monkey M, bottom: Monkey T. Difference in average angle between hyperplanes coding for the same variable (within) vs. coding for different variables (across). Negative values mean hyperplanes are better aligned when coding for the same variable. Dots: angular difference for data (dark gray) and permutations (light brown); dark gray connecting lines: significant results (p_val <0.05). c, Schematic illustration of the ‘state-space of subspaces’. Left: Hyperplane encoding color condition at time t1, represented as the LD1 vector of channel weights. This coding hyperplane rotates in time, and the evolution of the LD1 can be captured by reducing the dimensionality of the coding vectors. Right: Each hyperplane was reduced to two dimensions with PCA. A trajectory in the state-space of subspaces (PC1-PC2) describes hyperplane rotations over time, for activity (PC) and coding (LD) subspaces. d, Example trajectories in the state-space of subspaces. Top: trajectory of SEL cue information (LD1_color_, Mo180412002-pr1). Bottom left: trajectory of activity (PC1, Mo180412002-pr1). Bottom right: trajectory of SEL cue information (LD1_color_, Mo180619002-pr1); x-axis: trial time, y-axis: PC1, z-axis: PC2; color: ratio of weight strength between superficial and deep channels. e, Distribution of estimated oscillation periods for LD1_color_ across all sites showing significant periodicity (i.e. autocorrelogram peak above the 95^th^ percentile of permutations); x-axis: period (s), y-axis: number of sites. f, Autocorrelogram of the layer index for three example sites illustrating the three periodicity groups: sites with a period similar to LD1_color_, sites with a shorter period, and sites with no reliable periodic estimate; x-axis: time (seconds), y-axis: correlation, line: mean, shaded areas: 95% CI, color-code: periodicity type, red shaded area: chance level.

In Figure 7-a we focused only on selected reference coding subspaces at specific times, but linear discriminants can be computed at every time point along the task, and their fine laminar structure varies continuously (as seen in Figure 3). These laminar-weight variations can be interpreted geometrically (Figure 7-c) as a continuous change of orientation of the coding subspace encoding a specific variable of interest (e.g., selection cue color). Such dynamic reconfiguration can be described as a trajectory in the “space of subspaces”, mathematically a Grassmannian manifold over the space defined by laminar depths.

These trajectories of coding subspaces can be visualized using dimensionality-reduction approaches. For example, one can apply principal component analysis to the set of time-dependent linear discriminant axes associated with a given feature (e.g., LD1_color_) and project each time-resolved discriminant direction onto the first two principal components. Time can then be added as a third dimension, yielding plots such as Figure 7c (right), in which each point represents the LD1_color_ vector at a given time. The resulting trajectory captures the temporal evolution of the corresponding coding space.

In this representation, the color-coding subspace trajectory formed a characteristic helix, whose loops were paced by the task structure (Figure 7-d, top). This helical shape indicates that, after each visual cue, the principal axis of the coding subspace drifts along a specific trajectory that is approximately repeated across task blocks and reset to a similar starting condition at each new cue. Such helical trajectories were systematically observed across many sites, albeit with varying degrees of precision (Figure 7-d, bottom right; Suppl.Figure 6), and their estimated period matched closely the temporality of the task (Figure 7-e, significant autocorrelograms displayed in Suppl. Figure 8; visual cues were presented every 1.3s).

We emphasize that this peculiar organization of fine laminar-weight variations characterized the dynamics of coding subspaces, but not the raw MUA activity itself. At each time-point, the direction of maximum variability in neural activity is captured by the first principal component. Similarly to LDs, time-resolved PCs can be projected into a reduced space to describe the activity trajectory in the space of subspaces. However, this trajectory proved to be remarkably simple and largely unrelated to task structure (Figure 7d, bottom left, Suppl. Figure 1-a). Task-relevant information, by contrast, resides in smaller fluctuations along discriminant axes that are superimposed on this larger, unspecific variability. This is consistent with the observation that LD vectors show their strongest loadings on lower-rank PCs associated with smaller fractions of activity variance (Suppl. Figure 7). The instantaneous position of a discriminant axis thus indicates where to look for task-related information across laminar depths, and the trajectory of its evolution constitutes a trajectory of information-bearing fluctuations, tiny signals, composed of coordinated multilaminar activity, embedded within much larger, task-irrelevant variability. In short, while trajectories of raw activity fluctuations are trivial, trajectories of information-carrying modulations are highly structured.

To uncover potential spatial structure in the temporally organized variations of coding subspaces, we computed an index of layer involvement (see methods), such that large positive (negative) values indicate relative stronger weights in superficial (deep) layers. This index effectively captures the moving laminar “center of mass” of the coding subspace over time. When coloring the helical trajectories by this index, we observed a characteristic alternation of red and blue segments (Figure 7-d).

These alternations indicate that information-carrying fluctuations propagate through the cortical column, with the laminar center of mass shifting back and forth between superficial and deep layers. Similar dynamic patterns were observed across many sites (Supplementary Figure 6), although the specific mode of intra-columnar propagation was highly variable. In some sites, color-related information was initially concentrated in superficial layers and then propagated toward deeper ones (e.g., Figure 7d, top), whereas in others the pattern was reversed (e.g., Figure 7d, bottom right). Likewise, the temporal profile of information flow within the column was site-specific (example sites: Figure 7f; all periodic sites: Supplementary Figure 9). In other words, information cycled across laminar depths with variable periodicity: in more than half of the sites, the layer index displayed a clear periodicity, reflecting repeated shifts of information aligned with the dynamics of LD1 (Figure 7f, left). In some sites, however, this cyclic alternation occurred on a faster timescale than the full reshaping of LD1 (Figure 7f, middle), indicating that information could travel across layers more rapidly than the global reconfiguration of the coding subspace. Finally, a subset of sites showed no clear periodicity in layer index (Figure 7f, right), even when LD1 itself was periodic, pointing instead to a more irregular mixing of information across laminar depths.

Altogether, these results demonstrate that while propagating information across laminar depths is a ubiquitous feature of motor cortical dynamics, the direction, speed, and temporal pattern of the laminar “center of mass” are site-specific.

## Discussion

Our results show that task-related information is not confined to a few specific layers at any given time. Instead, it is encoded through the coordinated activity of the entire columnar circuit in the motor cortex. These coding subspaces -defined as lower-dimensional hyperplanes-are flexibly reused and recycled throughout the trial: related variables, such as cue identity and match, recruit overlapping subspaces, whereas the same variable, such as target direction, engages distinct subspaces depending on its behavioral context. Consequently, multiplexing of distinct task-related variables emerges from subtle inhomogeneities in layer contributions within highly distributed activity patterns. Moreover, this coordination-based encoding is dynamic, with alternating and time-varying enhancement of superficial and deep layer contributions.

### Cortical columns allow multiplexing in motor cortex

The extensive recurrent connections dominating the cortical columns allow them to transiently maintain incoming information and potentially operate as individual processing units **(Nolan *et al*., 2025)**. In this study, all laminar coding subspaces identified in the motor cortex were highly distributed, supporting the hypothesis that the functional unit corresponds to the entire column. In agreement with **(Opris *et al*., 2011, 2015; Opris, 2013)**, coordination across layers generated columnar activity patterns capable of encoding multiple task-related variables. In their case, neurons were modulated in both superficial and deep layers, reflected in our study as distributed lower-dimensional subspaces capturing the states and trajectories of the latent variables. In contrast, **Chandrasekaran et al. (2017)** argued for a layer-based computational unit in premotor cortex. They reported an overall superficial-to-deep decrease of visuomotor index, suggesting that decision-related activity was more prominent in superficial layers, together with an earlier onset of decision encoding in this layer (∼15ms). However, their depth profile of index followed a cubic trend, increasing again towards the deepest channel (at 2.5mm), and individual sites exhibited notable laminar variability. The difference of laminar coverage between their study (∼2.3mm) and ours (>4mm), along with this site-by-site variability, could therefore account for the apparent discrepancies between our results, especially considering that the reported cortical thickness of the motor cortex is around 3.5 mm **(Koo *et al*., 2012; Beul and Hilgetag, 2017)**.

If the cortical column operates as a distributed functional unit, a central question is how it can simultaneously encode multiple variables. At the single-unit level, multiplexing has often been understood through the framework of mixed selectivity, whereby neurons respond to combinations of task variables and thereby increase the dimensionality of population representations **(Rigotti *et al*., 2013; Fusi, Miller and Rigotti, 2016; Tye *et al*., 2024; Zhang *et al*., 2025)**. Our findings suggest that an analogous principle applies at the laminar scale: multiplexing arises from the coexistence of variable-specific subspaces within the cortical column. The partial overlap among these subspaces is reminiscent of neuronal mixed selectivity, raising the possibility that the column generates a high-dimensional laminar code capable of supporting the linear decoding of multiple task variables by distinct downstream readouts.

Despite layer V being traditionally considered the main output pathway of the motor cortex towards subcortical structures and spinal cord **(Jones and Wise, 1977; Dum and Strick, 1991; Harris and Shepherd, 2015)**, recent evidence indicates a more complex picture. For instance, **Borra et al. (2021)** reported equally strong cortico-striatal projections from layers III, V and VI, with distinct basal ganglia regions receiving characteristically weighted combinations of these laminar inputs. This organization suggests that downstream targets could, in principle, access parameter-specific laminar subspaces and exploit laminar multiplexing. Such subspace-specific communication channels, determined by source-target interactions, resemble the *communication subspace* concept **(Semedo *et al*., 2019)**, extended here to the columnar network. In this context, rather than involving only layer-to-layer interactions, the communication subspace may overlap with local coding subspaces, establishing instead a multi-layer-to-region flow of information.

### Different behavioral strategies across monkeys

In this study, monkeys learned a complex delayed match-to-sample task. Despite extensive training and overall correct behavioral performance, individual strategies may have emerged. The instruction was to remember the direction of the spatial cue matching the color of the selection cue. However, monkeys could either wait for the matching cue to appear or anticipate its occurrence, given that the order of the spatial cues was fixed. Based on the eye-movement profiles described for the two animals in **Nougaret et al. (2024)**, we believe that only monkey M anticipated the match. In that same study, we observed that uninstructed hand and eye movements were informative of color condition and were more stable in monkey T than in monkey M. This stability difference could account for the neural stability differences observed across monkeys (Figure 3-e), as well as for the sustained color-encoding space in monkey T (Suppl.Figure 2).

A further inter-subject difference concerned the lack of a consistent cue-identity subspace at SEL in Monkey T (Figure 4-f). Because trials were presented in blocks sharing the same selection-cue color (∼15 per block), the cue identity was already predictable before presentation, and thus may not have required transient encoding at SEL.

Thus, while inter-individual strategy differences might have shaped specific encoding features, the laminar and temporal organization of the coding spaces remained largely consistent across subjects

### Dynamic encoding of *cue* and *direction* in laminar subspaces

This task required the encoding of two independent parameters -cue identity and direction-whose behavioral relevance changed dynamically across epochs, providing a framework to investigate how laminar representations flexibly reorganize. The cue identity information (at SEL) had to be transformed into a match/non-match signal to gate spatial information at the valid SC. Direction, in turn, acted as the choice parameter, yet alternated between relevant and irrelevant contexts and had to be retrieved from working memory at GO.

The encoding of target direction during the preparatory delay preceding movement has been extensively characterized in motor cortex **(Tanji and Evarts, 1976; Wise, 1985; Riehle and Requin, 1989; Messier and Kalaska, 2000; Cisek and Kalaska, 2005; Churchland *et al*., 2006; Churchland, Santhanam and Shenoy, 2006; Thura and Cisek, 2014)**. Our results suggest that the population preparatory space is transiently visited during the delay following the informative cue, and reemerges during the final delay before GO (Suppl.Figure 5-b). The absence of sustained directionally selective preparatory activity in the motor cortex could be related to the long delay durations (e.g., more than 3s for blue trials) and the intervening distractor cues.

The directional tuning observed during these preparatory epochs typically differs from that during movement execution, with distinct subsets of neurons contributing to each **(Riehle and Requin, 1989)**, resulting in largely orthogonal subspaces in population state-space **(Churchland and Shenoy 2024)**. This dissociation between preparatory and executory subspaces parallels our finding that visual and motor subspaces accurately encoded direction, while the underlying coding subspaces were misaligned. Additionally, distractors were represented within the visual subspace -in one monkey-when presented before target information (Suppl. Figure 4). In the other cases, distractor directions were not represented within that subspace, possibly encoded along orthogonal directions, as suggested for the prefrontal cortex by **Qian et al. (2025)** and for the intraparietal sulcus by **Ritz and Shenhav (2024)**.

Beyond direction, the motor cortex is also known to represent abstract rules such as serial order **(Carpenter, Georgopoulos and Pellizzer, 1999; Carpenter *et al*., 2018)**. Since the cue color (at SEL) indicated order in the upcoming spatial-cue sequence, its encoding might reflect the temporal estimation of the match; consistently, this effect was only observed in the monkey anticipating the relevant cues. In addition, we found validity (match/non-match) encoding during the spatial cues -a higher-order, non-visual feature directly relevant for selecting the correct information.

Taken together, these results suggest that neural activity in the motor cortex organizes into lower-dimensional subspaces whose geometry reflects not only what is encoded, but also its behavioral relevance.

### Temporal recruitment of laminar coding subspaces

The idea that coordinated activity patterns across the cortical column define behaviorally relevant coding subspaces was further supported by their temporal reuse and recycling. Such subspace flexibility has recently been reported in the prefrontal cortex, suggesting that, beyond encoding specific variables, subspaces might be optimized for particular computations **(Wentz *et al*., 2025)**. Similarly, artificial neural networks trained on multiple tasks were shown to reuse neural representations and dynamical motifs **(Driscoll et al., 2024; Yang et al., 2019)**. Consistent with this view, monkeys performing compositional tasks exhibit shared neural subspaces across task contexts **(Tafazoli et al., 2026)**.

The low-dimensional structure describing the encoding of task-related variables is thought to form a behavior-specific neural manifold, containing all possible population states constrained both by intrinsic circuitry and extrinsic task demands **(Perich, Narain and Gallego, 2025)**. These low-dimensional trajectories capture only a small fraction of the total variability in the population (Suppl. Figure 7), consistent with previous reports **(Kobak et al., 2016)**. Nonetheless, the evolution of dynamics within these subspaces remains remarkably stable across neuronal populations and even across individuals of the same species **(Gallego *et al*., 2020; Safaie *et al*., 2023)**, as we observed when comparing target direction representations built from different cues (Suppl. Figure 3-a).

To continuously track these latent trajectories, coding subspaces themselves underwent a dynamic reconfiguration, describing a helical trajectory in the ‘space of subspaces’ (Figure 7-c, top). This temporal organization was shaped by task structure, with similar paths re-emerging after successive cues and thereby capturing the flow of color- and condition-specific information across the laminar circuit (Figure 7e). At the same time, the laminar weights defining each coding axis were continuously reshaped, causing the laminar “center of mass” of information to shift between superficial and deep layers in a site-specific manner. A tempting interpretation is that these helices reflect propagating waves **(Kuzmina, Kriukov and Lebedev, 2024).** In our case, however, they correspond not to waves of raw activity, but to waves of information. This distinction is important. The dominant low-dimensional trajectory of population activity obtained by PCA was simple, largely task-independent, and consistent with broad, homogeneous up- and down-scaling across layers (Figure 7c, bottom left). By contrast, the discriminant trajectories isolated weaker but behaviorally meaningful fluctuations embedded within this larger, nonspecific background. The helical paths traced by the coding subspaces are therefore more consistent with the progressive recruitment of distinct laminar subsets carrying task-relevant information than with a global wave of excitation sweeping uniformly across the column. In this sense, the column appears to host an information-bearing propagation front embedded within stronger, less specific “spontaneous” fluctuations. This interpretation is broadly consistent with proposals that transient propagation events support the spatiotemporal routing of cortical information **(Mohanta et al., 2024; Muller et al., 2018)**, here revisited in a multilaminar rather than horizontally spreading framework. In many sites, this propagation was task-locked, as indicated by the recurrence of similar trajectories across task epochs. In other cases, however, laminar alternations unfolded on a faster timescale than the full recurrence of the coding trajectory, indicating that information flow was governed not only by external task timing but also by intrinsic circuit dynamics **(Palkar et al., 2023)**. More irregular cases may reflect the interaction of multiple propagation fronts, whose combined effects could introduce faster components through interference or yield more complex, less periodic fluctuations of the laminar center of mass.

In conclusion, our results highlight a previously underappreciated principle: computation in motor cortex emerges from dynamic trajectories of multilaminar information flow, not from hardwired laminar specializations.

### Materials and Methods Experimental design Animal preparation

We analyzed an electrophysiological dataset collected in our laboratory and previously described in **Nougaret *et al*. (2024)**. In brief, the study included two adult macaque monkeys (M: 14Kg, T: 10kg, male, macaca mulatta). Animal care followed the European Commission Regulations (Directive 2010/63/EU on the protection of animals used for scientific purposes) applied to French laws (decision of the 1st of February 2013). All experiments took place in a licensed institution (B1301404), after being evaluated by the local Ethics Committee (CEEA 071) and approved by the French Ministry of High Education and Research (authorization 03383.02). The monkeys participating in this study were daily monitored to ensure their welfare and good physical condition. They were water-restricted in their cage, and liquid reward was provided during experimental sessions. If the intake during task performance was not sufficient (below 18ml/kg), extra water and/or fruit was given in their home cage. On weekends or resting days, they received the complete ration of liquid (water and/or fruits) in their home cage. Animals were pair-housed with free access to dry pellets and regularly provided with toys and enrichment. Data from the same two monkeys, but from the other hemisphere, was used in previous studies (**Confais et al., 2012a, 2012b, 2020; Kilavik et al., 2010, 2014**).

Monkeys were trained to perform a visuomotor delayed match-to-sample task (see details below). Once they had learnt the task, a cylindrical titanium recording chamber (19mm inner diameter) was implanted above the motor cortex (M1 and PMd), in the hemisphere contralateral to the trained arm. MRI scans prior to surgery as well as intra-cortical electrical microstimulation (ICMS; as described in **Asanuma and Rosén, 1972**) during the first recording weeks, ensured that multi-electrode recordings targeted upper limb regions (Figure 2-e).

### Behavioral setup and task

The behavioral paradigm, also described in **Nougaret *et al*. (2024)**, consisted of a highly temporally predictive delayed match-to-sample task (DMS) (Fig-2, a). The monkeys were presented with an initial color-coded selection cue (SEL), followed by three spatial cues (SCs) and a GO signal. The task required arm-reaching movements, after the GO, in the direction of the spatial cue matching the color of the SEL cue. A vertical computer monitor was used to display the visual scene, and monkeys used a handle freely movable in the horizontal plane to perform the movements. Visual targets appeared in the four possible diagonal directions.

A new trial was initiated when the monkey placed the cursor (small white square of 0.4cm edge) inside the central fixation dot (yellow flashing disc of 0.45cm radius). After a 50ms auditory tone and 1000ms central hand fixation, a selection cue (SEL) was displayed at the center of the screen during 300ms, indicating the color condition for the current trial. Specifically, SEL was one out of three possible colored polygons (blue, green or pink-3cm radius). The offset of SEL was followed by a 1000ms delay. After that, three spatial cues were sequentially presented (SC1, SC2 and SC3) for 300ms each, with fixed inter-cue delays of 1000ms. The three SCs corresponded to a sequence of three colored circles (0.9cm radius), blue-green-pink, always presented in the same order. For each SC, all four diagonal positions were equally likely, and independent of the position of the others; this implied that consecutive SCs could be presented in the same location. After a last 1000ms delay following SC3 (last spatial cue), the GO signal was presented. At the moment of the GO, the animal was required to perform the reaching movement in the direction of the memorized valid SC (i.e. matching the SEL color). The GO signal was not directionally informative. If the monkey touched the peripheral correct target within 500ms of reaction time, the trial was successfully completed and liquid reward was delivered.

All in all, there were 192 unique trials, given by 3 possible color conditions (color of SEL) and 4 independent locations for SC1, SC2 and SC3. Trials were presented in small blocks with the same color condition (∼15 trials), in order to facilitate the task. For Monkey T, only 3 out of 4 possible directions were presented each day (randomly selected in each behavioral session), to compensate for a less number of performed trials. Within the block, unique trials were pseudo-randomly selected, and incorrect trials were repeated at the end of the block until all trials were correctly executed.

Briefly, this behavioral paradigm corresponds to a temporally predictive delayed match to sample. The valid SC was cued at the trial start by SEL, and it could be temporally predicted (fixed order of SCs). The spatial location of the SCs (valid and invalid) was unpredictable.

### Data acquisition

During recording days (maximally 5 days a week), a multi-electrode, computer-controlled microdrive (MT-EPS, Alpha Omega, Nazareth Illith, Israel) was attached to the recording chamber and used to transdurally insert up to five single-tip microelectrodes (typical impedance 0.3-1.2MΩ at 1,000Hz; FHC) or up to two linear microelectrode arrays (either V-or S-probes, Plexon, Dallas, TX, USA or LMA, Alpha Omega; each with 24 or 32 contacts, inter-contact spacing either 100, 150 or 200µm; 12.5 or 15µm micrometer contact diameters) into motor cortex.

We define a ‘site’ as the neural recording obtained from a single electrode (single tip or linear array) within a given behavioral session. When multiple electrodes were simultaneously inserted, each recording was considered a separate site. In this study, we only considered sites recorded with linear arrays.

The arrays were positioned and lowered independently within the chamber (Flex-MT drive; Alpha Omega) in each session. Individual guide-tubes for each array were used that did not penetrate the dura (no guide was used for the more rigid LMA array). The reference was specific to each array type. For the LMA (Alpha Omega) it was an insulated wire exposed at the tip, either emerged in the chamber saline, or attached with a crocodile clip to the probe stainless steel tube (which in turn was lowered into the chamber liquid, but not extending into brain tissue, as the lower part of the probe was epoxy-insulated). For the V- and S-probes (Plexon) in most cases the reference was the stainless steel shaft of the array (extending into brain tissue, in near proximity to the probe’s recording contacts). In a few sessions, the reference was instead placed on a skull-screw on the more posterior headpost (6/36 sites using V-probes in monkey T) or on a screw on the saline-filled recording chamber (2/50 sites using S-probes in monkey M). For both array types, the ground was either connected to a skull-screw of the remote titanium head-fixation post, or to a screw of the titanium recording chamber.

All linear array recordings were obtained on a recording platform with components commercialized by Blackrock Neurotech (Salt Lake City, UT, USA). This system included Cereplex M digital head-stages (versions PN 6956, PN 9360 and PN 10129) connected to a Digital Hub (versions PN 6973, PN 6973 DEV 16-021, PN 10480) via custom HDMI cables (versions PN 8083, PN 8068), which transmitted signals via fiber optics to a 128 channel Neural Signal Processor (NSP hardware version 1.0), and control software Cerebus Central Suite (v6.03 and v6.05 for monkeys T and M, respectively; running on Windows 7). Neuronal signals were hardware filtered (0.3Hz – 7.5 kHz), digitized and saved for offline analysis at a sampling rate of 30 kHz. Behavioral event codes (TTL, 8 bits) were transmitted online to the DAQ system from the VCortex software (version 2.2 running on Win XP; NIMH, http://dally.nimh.nih.gov), which was used to control the behavioral task.

### Linear probe insertion and cortical layer estimation

For each site, while lowering the linear probes, the neuronal activity was monitored to determine the depth at which the first neuronal signals could be heard (hash) and seen on DAQ monitor, on the deepest probe channels. Typically, the contacts just above the ones with the first multiunit activity would have a clear heartbeat signal in the LFP, confirming their position inside the dura. The contacts above would typically have much more noise (incl. 50Hz line noise), since we only filled the recording chamber with saline after having correctly positioned the probe. The probes were then moved down in small regular steps of 50-100 microns (micrometers), while observing that the neuronal and heartbeat signal would shift upwards along the contacts. When all but 2-3 contacts were inside the brain, the probe was left in place for 1-3H, to allow the tissue to settle around the probe before we started the neuronal recordings. If we then observed neuronal activity on all contacts, we retracted the probe until we observed heartbeat on the top 1-2 contacts, and neuronal signals only from contacts 2 or 3 and below. Sometimes the neuronal tissue would still drift up slightly around the probe during the recordings, but typically less than the spacing between neighbouring channels. To counteract the effect of such drift in the MUA signal, we adopted a drift correction pre-processing step (see below).

A slightly tilted adapter added to the recording chamber ensured that the probes would penetrate the cortex perpendicular to the cortical layers, as long as we remained sufficiently distant from landmarks such as the central and arcuate sulci and the precentral dimple. The positions of these landmarks were estimated from anatomical MRI scans and confirmed with the ICMS chamber mapping done with single electrodes in the initial recording sessions.

Cortical layers were defined based on depth from the dura, individually for each site. The macaque M1 and PMd is about 3.5mm thick, on the gyral surface **(Koo *et al*., 2012; Beul and Hilgetag, 2017)**. This thickness was confirmed in our animals with anatomical MRI). As the macaque cortex has an approximate equal thickness of superficial and deep layers **(Hutsler, Lee and Porter, 2005)**, we considered the upper 1.8 mm as superficial layers (L2/3) and anything below as deep layers (L5/6). Layers 2 and 3 were grouped together -given the absence of a clear anatomical boundary- and instead subdivided into L23-SUP (superficial half) and L23-DEEP (deep half). Channels extending 0.85 below L23-DEEP were assigned to layer 5 (L5), and deeper channels to layer 6 (L6). Since some linear arrays spanned up to 4 mm depth, L6 channels may have included interstitial neurons from the underlying white matter. These interstitial neurons are thought to represent late-born layer VI neurons that failed to fully migrate into the cortical plate **(Kostovic and Rakic, 1980)**. Given their shared morphological and functional properties, they are often grouped with L6 neurons.

Measurements like current source density (CSD), extensively used in primary visual cortex to estimate layer IV, are less reliable in higher order cortical areas with a less clearly defined layer structure, like motor/premotor cortex, notably regarding the input L4.

## Data analysis

### Neural dataset

As mentioned above, in this study we restricted our analysis to neural data recorded with linear electrode arrays, which allowed us to capture population activity across the full cortical column simultaneously. Since laminar probes were acutely inserted at different cortical locations, achieving perfect channel-to-channel vertical alignment across sites was challenging. Therefore, each laminar site was treated as an independent dataset in all analyses. To compensate for the reduction in statistical power, we included only behavioral sessions in which the monkeys completed at least 90 correct trials (corresponding to approximately two entire blocks of all color conditions, with ∼15 trials per block).

The resulting dataset contained 27 sites for monkey M, and 14 sites for monkey T, obtained across 19 and 10 behavioral sessions, respectively. All sites included channels spanning at least three distinct cortical layers, and were distributed along the PMd/M1 cortical sheet. Both animals had successfully learnt the task, as indicated by an average performance of 80% of correct choices in complete trials (hand fixation maintained until the GO), as reported in **Nougaret *et al*. (2024)**.

### Preprocessing

Extracting the Multi-Unit Activity (MUA)

To obtain a neural population measure, we extracted the multi-unit activity (MUA) as the envelope around the high-pass filtered continuous signal, as described in **Stark and Abeles (2007)**. The step-by-step process involved:

- Filter the raw 30 KHz electrophysiology continuous signal from each channel using a band-pass filter (300-6000Hz).
- Clip the values above and beyond +/-2 standard deviations.
- Rectify the signal by squaring it.
- Smooth the continuous data by low pass filtering it at 250Hz, and downsample it to 1 KHz.
- Negative values resulting from filtering were set to zero, and take the square root of the signal.

The MUA represents the superimposed activity of multiple neurons around the electrode. It reflects the population signal even in the absence of sortable single units, allowing us to sample across all laminar channels. Furthermore, it captures the local activity in the vicinity of each contact, respecting the laminar differences on the underlying neuronal populations.

### Correcting for spatial drift

As mentioned above, because the laminar probes were acutely inserted, spatial drift occurred during some sessions, even if we waited after inserting the probes before initiating the recordings. This drift was reflected as slow baseline shifts in the MUA signal and by the appearance or disappearance of single units over time. After confirming that the time-scale of the drift was much longer than any task-related fluctuations, we corrected it by subtracting a moving average from the MUA signal. This moving average, serving as an ‘online baseline,’ was computed by smoothing the signal with a 30s window using *uniform_filter1d* (from *scipy.ndimage*). The 30s timescale was chosen to be longer than individual trials but shorter than a condition block, ensuring that non-artifactual fluctuations in neuronal activity were preserved. The correction was applied independently to each channel, and edge effects were discarded. In order to center data around zero, the corrected MUA in each channel was z-scored.

### Alignment between the neuronal signal and the behavior

We here only analyzed behaviorally correct trials. In most of the analysis presented in this manuscript, the MUA was aligned to the initial selection cue (SEL). Complete trials were defined from 500ms before the monkeys touched the central target (corresponding to 1s before SEL) until the average target-touch time for that session. Analysis studying the preparatory period of the movement were instead aligned on the GO signal. In general, since all temporal delays were stable across conditions, aligning trials at the start (SEL) strongly preserved their temporal synchronization.

### Removal of artefactual channels

Channels with extremely bursty neurons (detected using offline spike sorting) were excluded to avoid biasing results toward a single persistent-firing neuron. This was rare: only 4 channels (<<0.5% of all sites) were removed.

### Single-channel temporal evolution

Single-channel MUA activity was grouped by color condition (SEL identity) or movement direction and then averaged across trials. The shaded area represents the 95 confidence interval of the mean. A Savitzky-Golay filter of 5th order with 200ms window was applied for temporal smoothing, using the *savgol_filter* function (from *scipy.signal*). To study the temporal modulations associated with movement, the trials were grouped by color condition, since the movement information was given at different times for the three different color conditions.

### Time-resolved dimensionality reduction

Two dimensionality reduction methods were used throughout this manuscript: Principal Component Analysis (PCA) and Linear Discriminant Analysis (LDA), implemented using the *sklearn.decomposition* and *sklearn.discriminant_analysis* modules, respectively. Both methods generate low-dimensional representations of high-dimensional data, in this case, using linear combination of channel weights. To avoid overfitting, models were trained on a random subset of trials (50% for color, 70% for direction) and tested on the remaining trials. Overlapping windows of 200ms, sliding by 25ms, were used to estimate the model iteratively across time.

- *PCA* captures the direction of maximum variability of the data, and it was applied to single-trial activity in an unsupervised manner.
- *LDA* finds the axes that best split the data in two or more given classes. It is a supervised method which works at the single-trial level, by providing to the model both the neural data (in all channels) and the categorical variable (color condition or movement direction).

After dimensionality reduction, *weights* and *projections* are extracted. The weights represent the contribution of each of the channels in the laminar probe, and the projections the specific state of the population activity in the lower-dimensional space.

Because PCA/LDA have sign indeterminacy, the orientation of the axes can flip when recomputing the model iteratively in different windows along the trial, leading to mismatches across time. To address this, we enforced a consistent sign by selecting the orientation that maximized projection stability, and adjusted both weights and projections accordingly. The temporal evolution of LDA/PCA projections is generated by concatenating the individual-window projections, assigning them the temporal value of the middle of the window. Weights were visualized as vertically arranged heatmaps, aligned with probe orientation, in which the color-code represented the contribution of each channel (e.g., Figure 3).

### Mutual information

We used mutual information (MI), specifically *WfMi* from *frites.workflow* (Combrisson et al., 2022), to quantify the amount of information about a behavioral variable contained in the single trial data, at individual sites. The numerical values are normalized by dividing by the theoretical maximum MI, given by log_2_(#*categories*). Only significant values of MI were plotted (computed using permutation test).

### Entropy

We used Shannon entropy (*H*) to quantify how distributed or localized the vector weights in each site were. To compute this, the weight vector was first normalized, squared, and converted into a probability distribution. Entropy was then computed as:

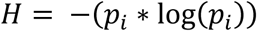

where *p_i_*denotes the normalized squared weight of channel *i*.

A low entropy value indicates that the vector is dominated by a few channels (localized), whereas a high entropy value indicates that contributions are more evenly distributed across channels. To account for the varying number of probe channels across recording sites, we normalized the entropy by its maximum possible value, log(*NN*), where *N* is the number of channels. After normalization, an entropy of 1 indicates equal contributions from all channels, while 0 indicates that only a single channel contributes.

### Similarity

We quantified the temporal stability across windows of the weight vectors per site using the absolute cosine similarity:

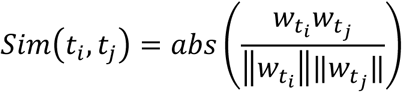

where *w*_*ti*_ and *w*_*tj*_ are the weight vectors at times *c*_*i*_ and *c*_*j*_, respectively. Taking the absolute value compensates for the sign indeterminacy inherent in PCA/LDA solutions. Similarity values range from 0 (orthogonal vectors) to 1 (perfectly aligned vectors).

To assess the consistency of the results, the weight vectors were recomputed across 50 bootstrap resamples of trials, cosine similarity was evaluated on these resampled vectors, and the final similarity for each site was obtained as the average across bootstraps.

### Subspace 2D analysis

Population activity in a fixed temporal window around the event of interest was used to fit an LDA model. Neural activity at other times was then projected into the same 2D subspace. Trajectories were temporally smoothed and condition-averaged for visualization purposes (both train and test trials). All quantitative analyses were performed on single-trial MUA.

### Cue identity space (at SEL)

#### Building the LDA space

To define the cue identity space, the LDA was trained to separate the three cue colors presented at SEL (blue, green, pink). A 300ms window starting 150ms after SEL onset was used. Half of the trials (50%) were randomly selected for training, ensuring class balance. To increase the number of training samples, each trial was subdivided into non-overlapping 50ms bins, resulting in six samples per trial. Remaining trials were used for testing, averaging each test trial over the 300ms window around SEL to obtain a single projection in the 2D space.

#### Generalization of color space

To test whether the cue identity space generated at SEL also encoded color at later time points, single-trial activity at the spatial cues (SCs) was projected into the 2D axes defined at SEL, without retraining the LDA model. For evaluation, trial labels were assigned according to the color presented at the SC: SC1 = blue, SC2 = green, SC3 = pink. A 300ms window (onset +150ms) was used for the projections.

#### Finding the validity axis

To define a validity axis within the cue identity space, single-trial activity around the SCs was projected into the SEL-defined 2D axes. Training trials were re-labeled as valid or invalid based on the color match between SEL and SC (e.g., for SC1: blue trials = valid, green and pink = invalid). Using these re-labeled training trials, a new LDA was trained on the 2D coordinates of the SEL space to separate valid from invalid cues. Test trials were projected onto this validity axis to evaluate generalization. Chance level was estimated by recomputing the validity axis with shuffled validity labels.

### Visual target direction space (at SCs)

#### Building the LDA space

To define the visual target direction space, LDA was trained to separate the directions (three for monkey T or four for monkey M; see above) presented at a given SC. A 300ms window starting 150ms after SC onset was used. Since the informative SC depended on the color condition (SEL color), only matching trials were used at each SC. 70% of these matching trials were randomly selected for training, ensuring class balance. To increase the number of training samples, each trial was subdivided into non-overlapping 50ms bins, resulting in six samples per trial. Remaining trials were used for testing, averaging each test trial over the 300ms window around the matching SC to obtain a single projection in the 2D space.

#### Generalization at other cues

To test whether the target space generalized across SCs, test trials from other color conditions at their respective matching SCs were projected into the LDA space defined above. For example, a space trained to separate target directions in blue trials at SC1 was used to project green trials at SC2, to assess whether the directional code was maintained. A 300ms window starting 150ms after matching SC was used for the projections.

#### Encoding distractors

To evaluate encoding of distractor directions (irrelevant SCs), non-matching trials were projected into the same LDA space. Using the same example, a space trained to separate (valid) target directions in blue trials at SC1 was used to project green trials at SC1, to assess whether the (blue) distractor direction was represented.

#### Generalization as a preparatory space

To determine whether the LDA space encoding visual target direction at SCs corresponded to the directionally-tuned preparatory activity shortly before movement onset, single-trial activity was projected into the LDA space using a 200 ms window aligned to the GO signal.

### Single-trial distance metric

All 2D projections were analyzed at the single trial level. For each trial, the Euclidean distance between the projected point and each class center (computed as the class-average of the training data) was calculated. The distance difference was defined as:

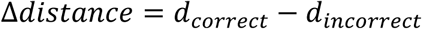

where *d_correct_* and *d_incorrected_* are the distances to the correct and incorrect class centers, respectively. If multiple incorrect classes existed, their distances were averaged.

The distance difference (Δ*distance*) was computed across all test trials for each train/test split. Values close to zero indicate no information about the class, whereas negative values indicate that trials are closer to their correct class.

To assess robustness, the entire procedure (including recomputing the LDA model) was repeated over 200 bootstrap resamples. At each bootstrap, five additional shuffles of class labels were performed to estimate chance-level performance.

### Significance testing

Significance was assessed using permutation testing to estimate the chance distribution for each analysis. In general, a measure (in most of the cases the mean distance difference) was considered significant if it fell below the 5th percentile of the permutation distribution.

P-values were computed based on the overlap between the empirical and chance distributions. When multiple comparisons were performed simultaneously, p-values were adjusted using Bonferroni correction.

### Angle between subspaces

To quantify the alignment of coding subspaces in the high-dimensional space, we computed the angles between the hyperplanes defined by LDA weight vectors, following the approach in **Knyazev and Argentati (2002)**. The *subspace_angles* function from *scipy.linalg* was used to calculate these angles across the different 2D planes. Since each pair of 2D planes yields two principal angles, we took the average of these two values to obtain a single measure of alignment per plane pair. Larger angles indicate stronger misalignment, whereas smaller angles indicate greater similarity between subspaces.

### Trajectory in the Grassmannian space

To examine the temporal evolution of the LDA axes in the space of subspaces, laminar weights vectors were reduced to two dimensions using PCA. Absolute values of the weights were used to correct for sign indeterminacy. The resulting trajectories were color-coded according to a layer index, indicating whether superficial or deep layers contributed more strongly at each time point.

### Index of layer involvement

To quantify the relative contribution of superficial and deep cortical layers to each LDA axis, we computed an index of layer involvement. First, we took the absolute value of the LDA weights and computed the average contribution across the weights in each layer, normalized by the sum of layer-averaged contributions, so that the total contribution summed to 1. We then averaged the contributions within superficial layers (L23-SUP, L23-DEEP) and deep layers (L5, L6). Finally, the layer involvement index was computed as the ratio of superficial to deep contributions. An index of 1 indicates equal contribution from superficial and deep layers, values greater than 1 indicate stronger superficial involvement, and values less than 1 indicate stronger deep involvement.

### Variance explained by LDA

Because LDA maximizes inter-class separability rather than total variance, it does not directly provide the amount of variance captured by each discriminant axis. To estimate the variance along each LD axis, we computed its alignment with all principal components (PCs). The variance of each LD axis was then obtained as the weighted sum of the variance explained by each PC, with weights given by the squared alignment between the LD axis and each principal component:

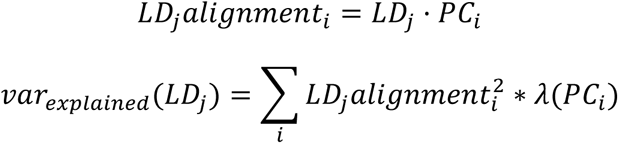

Where *i* corresponds to the index of the PC, *j* to the index of the LD and λ to the eigenvalue of the PC component, representing its variance explained.

### Periodicity of LDA_color_

To assess the temporal periodicity of the LD1_color_, we used the similarity matrix for each recording site. Starting at SC1, we extracted the local similarity profile as *S*(*t* − *τ*, *t* + *τ*), with a lag of τ ±2s. Then, the profile was z-scored and we computed its autocorrelation. We extracted similarity profiles -and their respective autocorrelation-along the diagonal, until the ± τ reached the end of the trial. All autocorrelations were then averaged to obtain a site-level estimate of temporal structure.

In parallel, we assessed the statistical significance of the autocorrelation profile. We generated surrogate distributions by shuffling the similarity profiles in blocks, with block lengths uniformly sampled between 0.6 and 1s, preserving local smoothness while disrupting long-range temporal organization. For each lag, the 5th–95th percentile range of the surrogate autocorrelation was computed. A site was considered to exhibit significant periodicity if the first non-zero-lag peak of the empirical autocorrelation exceeded the 95th percentile of the surrogate distribution. The lag of this peak was taken as the estimated period.

### Periodicity of the layer index

To assess the temporal periodicity of the layer index, we first extracted ±2s windows around randomly sampled reference time points after SC1 and computed the normalized autocorrelation of the layer index within each window. We repeated this procedure across 10.000 bootstraps with replacement. We obtained the mean and the confidence interval of the periodicity (5^th^-95^th^ percentile). In parallel, we assessed the statistical significance of the autocorrelation profile. We generated block-shuffled surrogate data (block lengths uniformly sampled between 0.3 and 0.8s), preserving local smoothness while disrupting long-range temporal organization. A site was considered to exhibit significant periodicity if a non-zero-lag peak (specifically the 5^th^ percentile across bootstraps) of the empirical autocorrelation exceeded the 95th percentile of the surrogate distribution. The lag of this peak was taken as the estimated period.

## Acknowledgments

The authors thank Nicolas Meirhaeghe for valuable scientific discussions during the preparation of this work; Joel Baurberg, Xavier Degiovanni and Luc Renaud for technical assistance; Sébastien Barniaud, Laurence Boes, Frédéric Charlin and Marc Martin for animal care.

## Funding

EU Horizon 2020 Skłodowska-Curie Actions grant, In2PrimateBrains – 956669. FLAG-ERA grant PrimCorNet, ANR-19-HBPR-0005. ANR FunSy grant, ANR-25-CE45-2766. Neuroschool end-of-phd grant from the French government under the “France 2030” investment plan managed by the French National Research Agency and from Excellence Initiative of AixMarseille University.

## Declaration of interests

The authors declare no competing interests.

## Supplementary Figures

**Suppl.Figure 1.**
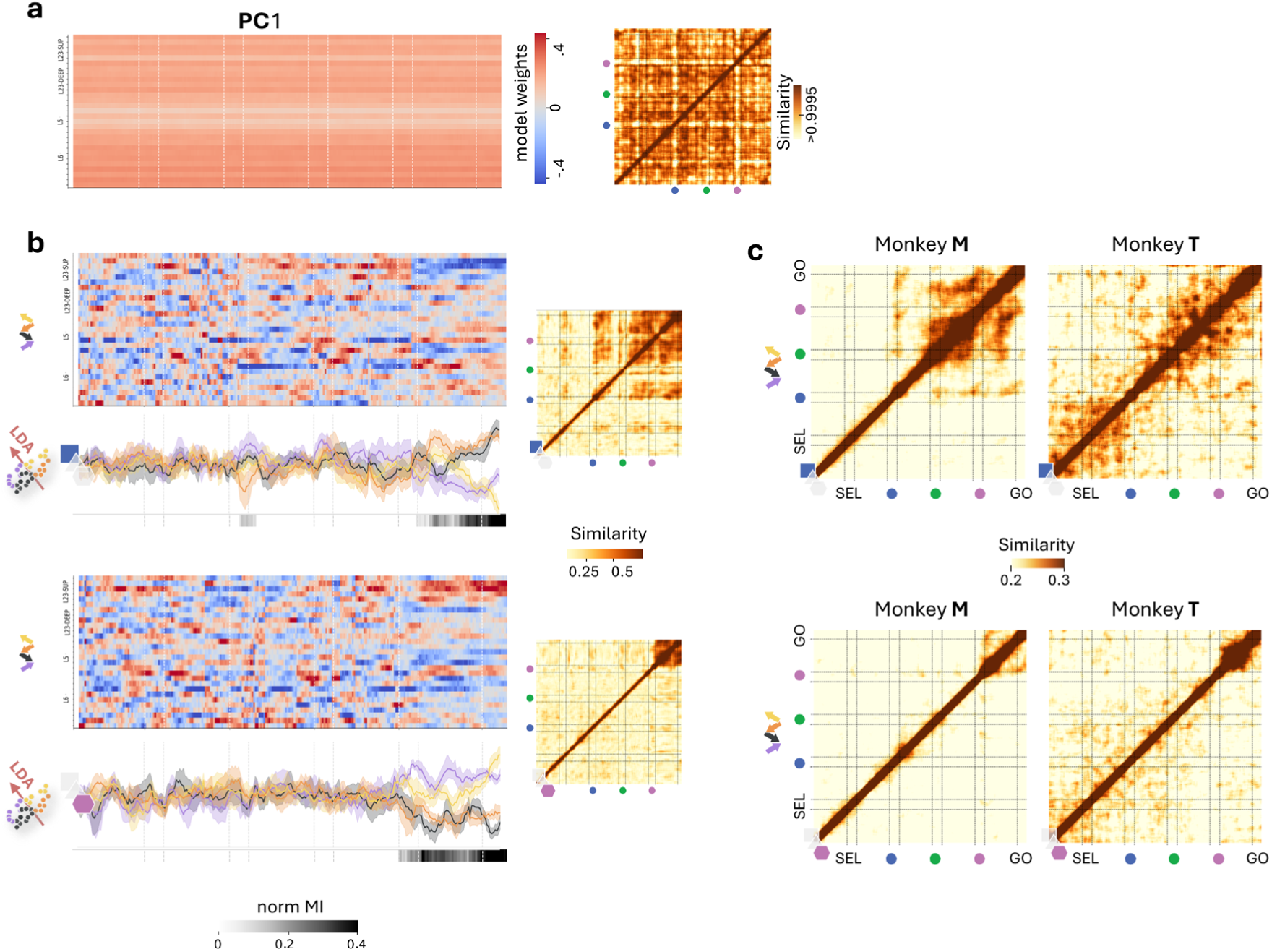
Activity spaces evolution (PCA) and direction coding spaces (LDA_direction_) for blue and pink trials. a, Left: Heatmap of weights from time-resolved PCA models for one example site. Right: Cross-temporal map of absolute cosine similarity across pairs of weights shown on the left. b, Top panels: Heatmaps displaying the weights of all time-resolved LDA_direction_ models for the example site using blue (top) and pink (bottom) trials. Bottom panels: Neural activity projected onto the first axis of LDA_direction_ (top: blue trials; bottom: pink trials), computed in overlapping 200ms windows and concatenated across the trial. Condition means (movement direction) are displayed as color-coded lines with 95% CI (shaded). Dark bars below indicate significant mutual information (MI), with intensity reflecting magnitude; white indicates non-significant bins. Right panels: cross-temporal map of absolute cosine similarity across pairs of weights in LDA_direction_, for blue (top) and pink (bottom) trials. Each map is the average of 50 similarity matrices from resampled training data. c, Same as in (b, right) but averaged per monkey across all sites; left: Monkey M, right: Monkey T, top: blue trials, bottom: pink trials. (a-b), Example site: Mo180412002-pr1.

**Suppl.Figure 2.**
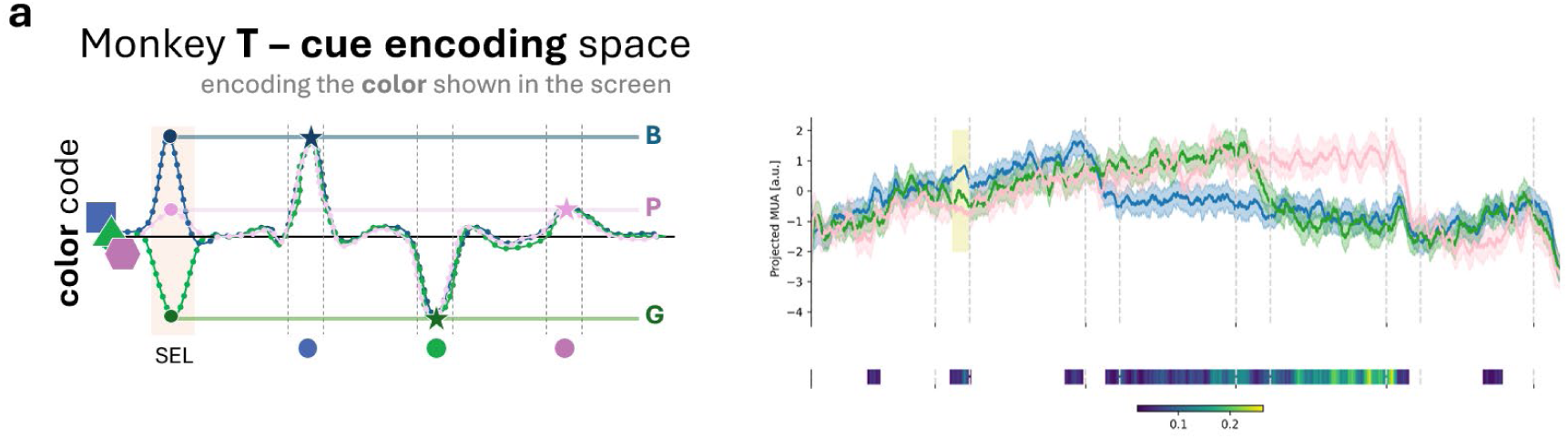
Sustained color-condition space in Monkey. **T.** Left: Schematic illustration of reusing the cue identity space at SEL to encode cue color transiently at SC1 (blue), SC2 (green) and SC3 (pink). Right: Example site. Trial-averaged projections on LD1_color_, separated by SEL color; line: mean, shaded: 95% CI, x-axis: trial time. Color-bar below indicate significant mutual information (MI); white indicates non-significant bins.

**Suppl.Figure 3.**
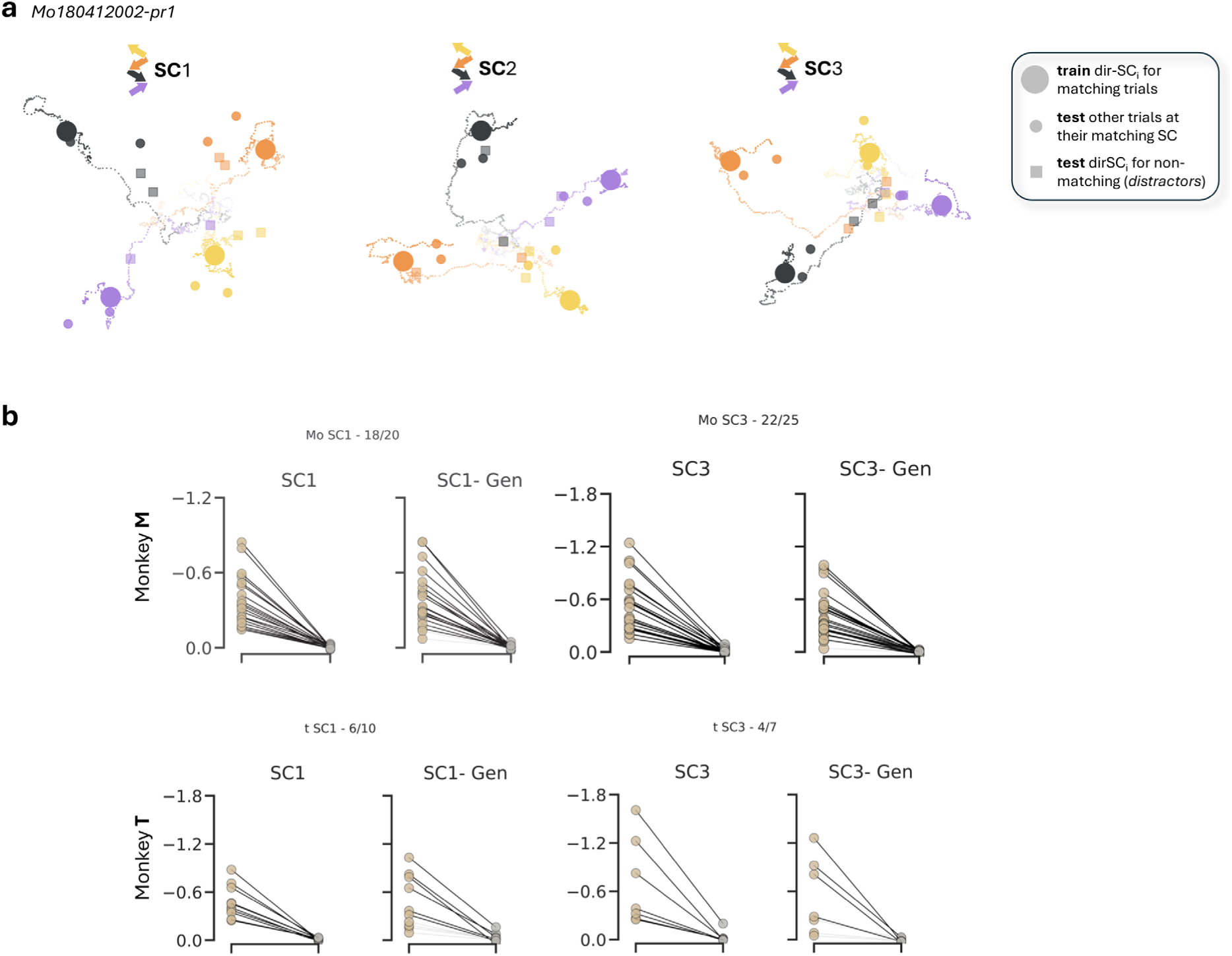
Generalization from other SCs. a, Example site. Condition-averaged trajectories around the training epoch when models were built at SC1 (left), SC2 (middle) and SC3 (right) for target direction on matching trials. Large colored dots represent the state of maximum cross-condition variance during the train epoch (matching trials). Small color-coded dots represent trial-averaged projections for correct trials at their respective matching SC. Small color-coded squares represent trial-averaged projections for non-matching trials, i.e., trials projected at the train epoch but coded by the direction of the presented cue. Color code: yellow: lower left, orange: upper left, purple: lower right, black: upper right. b, Summary of the single-trial distance metric (Δ_dist_) across sites for models trained at SC1 (blue trials, left) and SC3 (pink trials, right). Only sites with a significant SC target direction space (Δ_dist_ data < 95^th^ percentile of permutations) were tested for generalization. Dots: mean Δ_dist_ for each test: SEL, H_1_ (data: brown, permutations: gray); dark connecting lines: significant results after Bonferroni correction. Top: Monkey M; bottom: Monkey T.

**Suppl.Figure 4.**
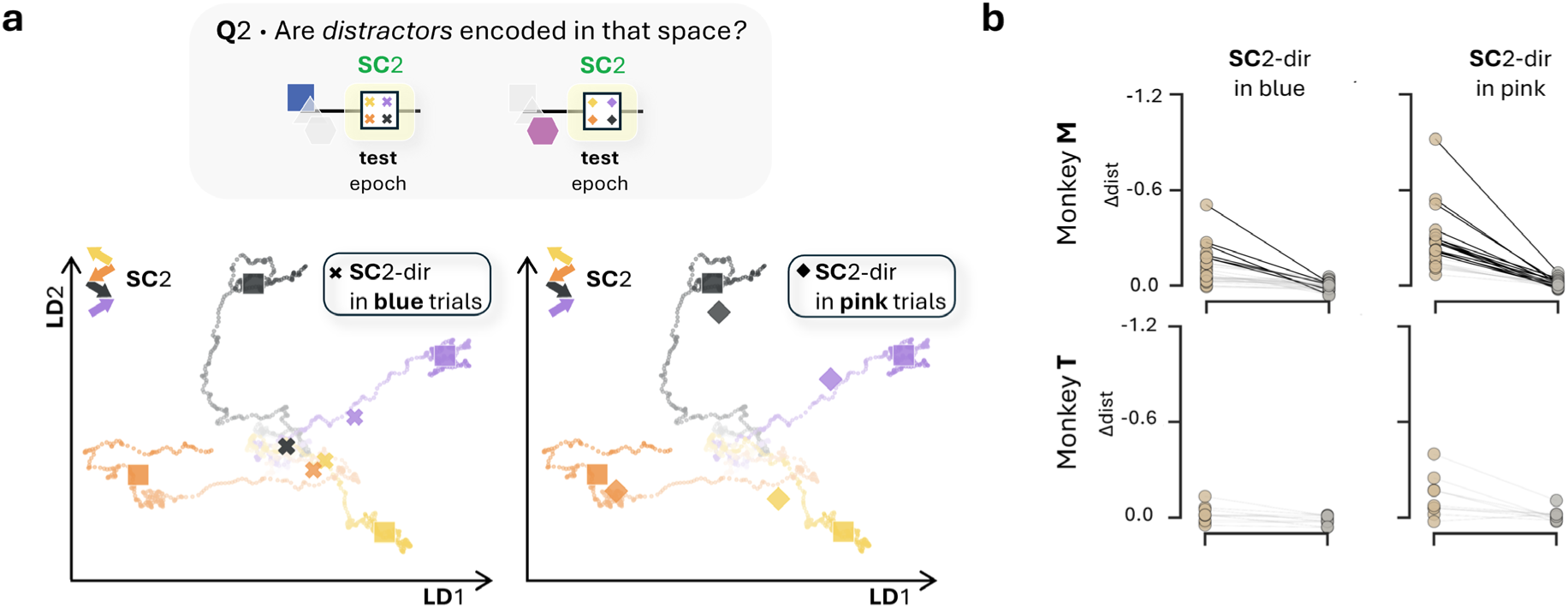
Encoding distractors. a, Top: Schematic illustration of distractor encoding. In blue (left) and pink (right) trials, the direction of SC2 (green) acted as a distractor rather than a target direction. We tested whether the distractor direction-state (SC2), i.e. direction of SC2 for non-matching trials, overlapped with target direction states (directional split of green trials at SC2). Bottom: Example site. Condition-averaged trajectories for blue (left) and pink (right) trials projected onto the SC2 target direction space (built on green trials). Color code: SC2 direction; Cross: average of blue trials at SC2, diamond: average of pink trials at SC2. b, Summary of the single-trial distance metric (Δ_dist_) across sites for models trained at SC2 (green trials) and tested in the direction of SC2 in blue (left) and pink (right) projections. Only sites with a significant SC2 target direction space (Δ_dist_ data < 95^th^ percentile of permutations) were included. Dots: mean Δ_dist_ for each test (SC2 direction in blue and pink trials; data: brown, permutations: gray). Dark connecting lines: significant results after Bonferroni correction. Top: Monkey M; bottom: Monkey T.

**Suppl.Figure 5.**
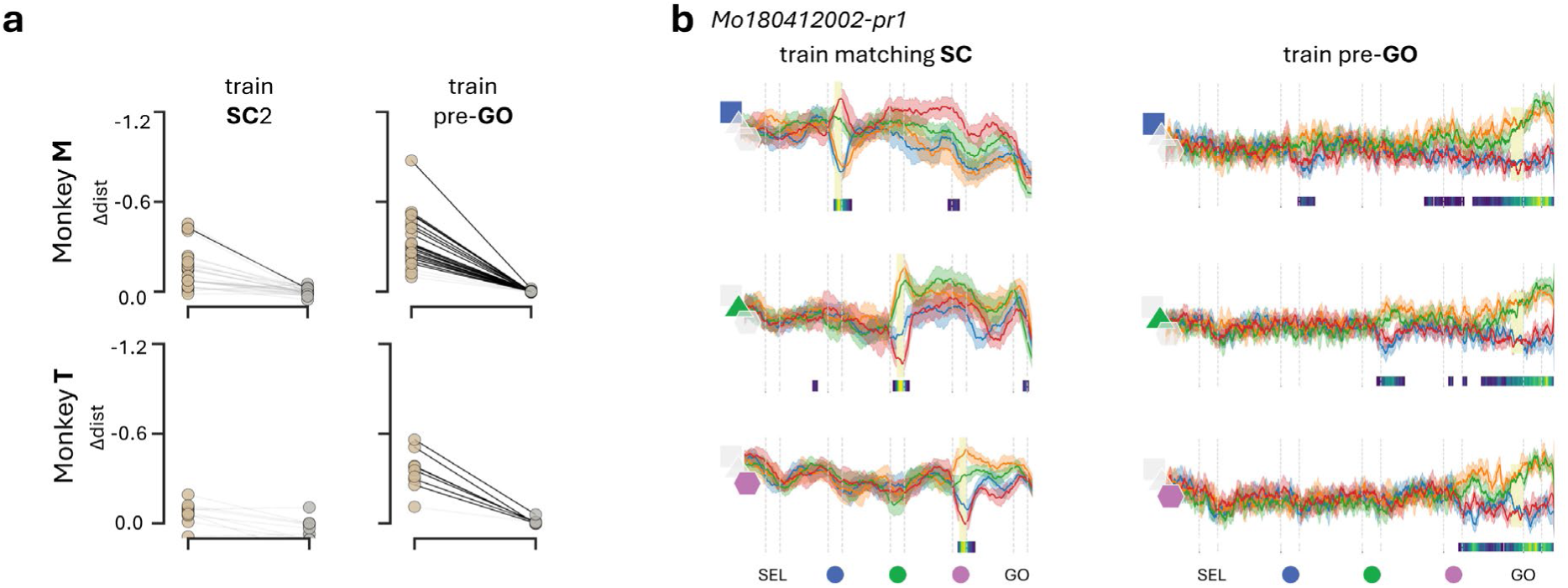
Cue-direction space vs Preparatory space. a, Summary of the single-trial distance metric (Δ_dist_) across sites for generalization of the cue space to the preparatory epoch (left) and existence of preparatory space (right). Only sites with a significant SC2 target direction space (Δ_dist_ data < 95^th^ percentile of permutations) were included for SC2 generalization (left). Top: Monkey M, bottom: Monkey T. Dots: mean Δ_dist_ for each test (data: brown, permutations: gray); dark connecting lines: significant results after Bonferroni correction. b, Example site: Mo180412002-pr1. Neural projections on an LDA_direction_ model built during matching cues (left) or in the preparatory epoch (pre-GO, right), separated by color condition (top: blue, middle: green, bottom: pink). Bottom colored-bar represents the time-resolved mutual information; white indicates non-significant bins.

**Suppl.Figure 6.**
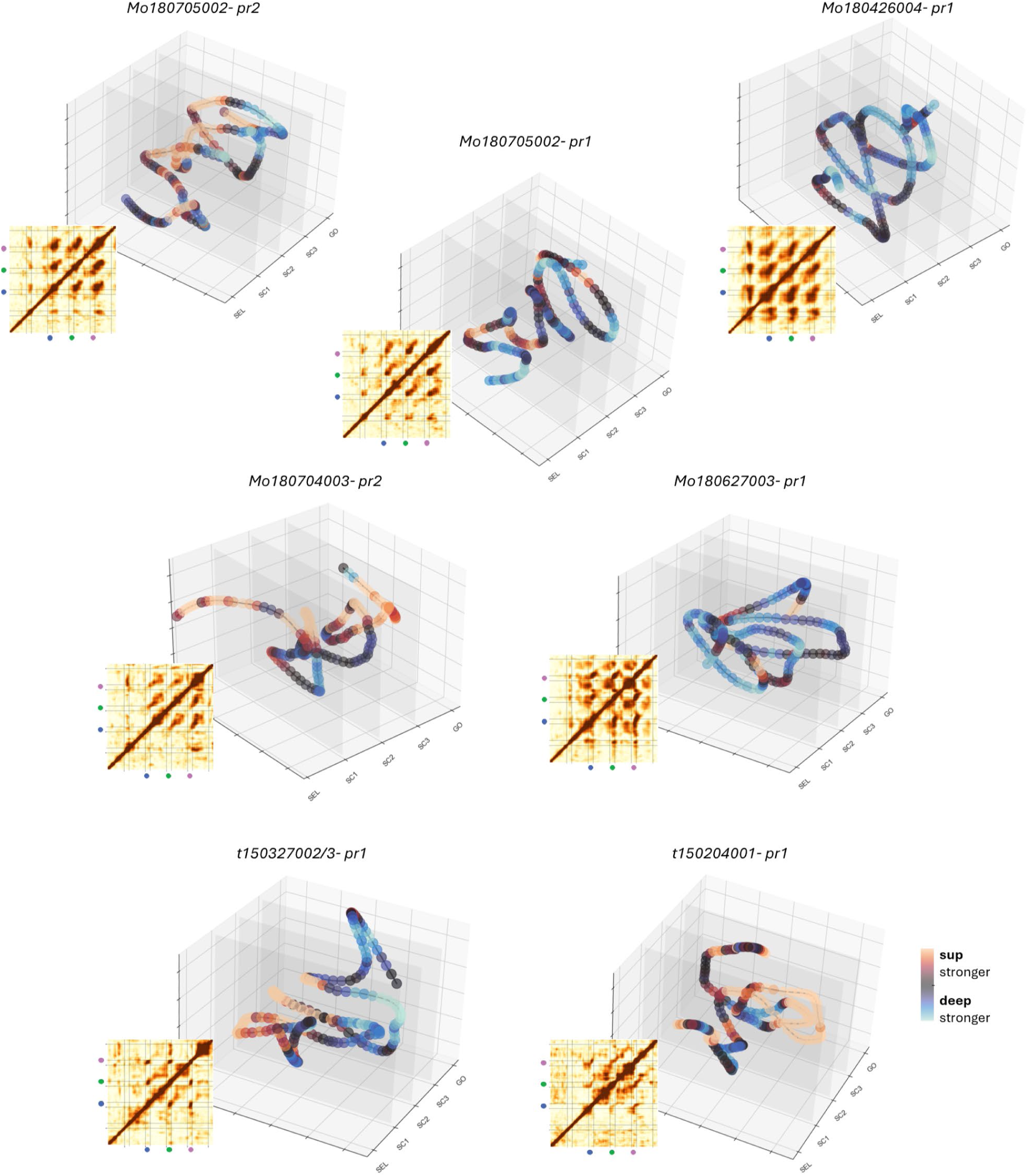
Trajectory of the coding vectors in the space of subspaces. Sites displaying a clear off-diagonal stripe in the similarity matrix are displayed. Right for each site: Site-specific trajectories of cue information (LDA_color_) in the state-space of subspaces; x-axis: trial time, y-axis: PC1, z-axis: PC2; color: ratio of weight strength between superficial and deep channels. Left for each site: Cross-temporal map of absolute cosine similarity across pairs of weights in LD1_color_. Each map is the average of 50 similarity matrices from resampled training data in each site; x-axis, y-axis: trial time.

**Suppl.Figure 7.**
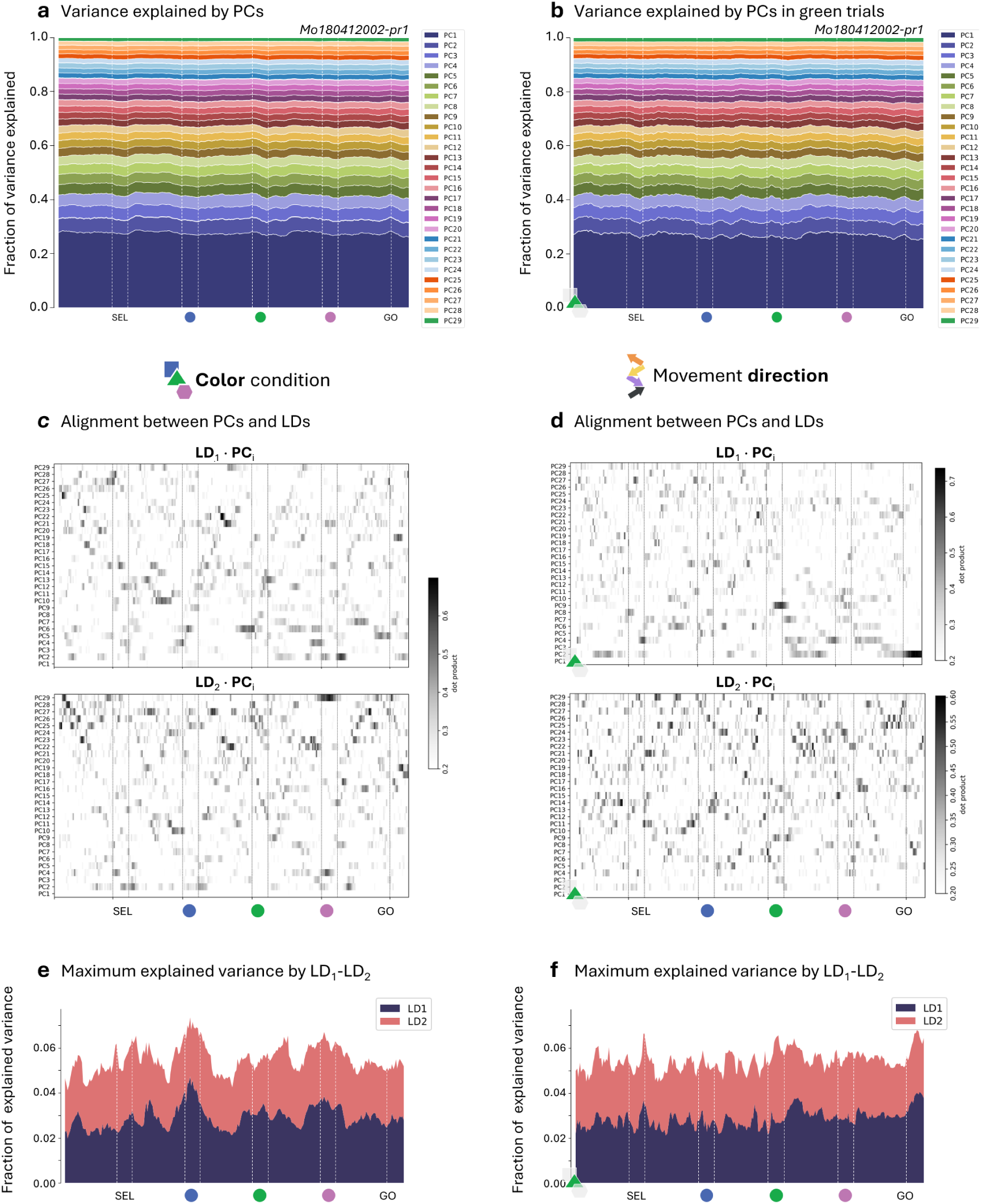
Variance explained by time-resolved PCA and LDA. a, Relative variance explained by each principal component, computed across all single trials at a representative site; x-axis: time, y-axis: fraction of variance explained, color: PC_i_. b, Same as a, but PCA computed using only on green trials. c, Alignment between LD_color_ and PCA (using the PCA in a). Time-resolved dot product between LD_1_/LD_2_ and each principal component; x-axis: time, y-axis: PC index, color: absolute dot product value. d, Same as c, but for LD_movement_ computed on green trials (using the PCA from b). e, Maximum cumulative variance explained by LD1 and LD2 (color-coding). At each time-point, this was computed as the weighted sum of the variance explained by each principal component, with weights given by the squared alignment between the component and LD1 (or LD2).; x-axis: time, y-axis: fraction of explained variance, color: LD index. f, Same as e, but for movement direction.

**Suppl.Figure 8.**
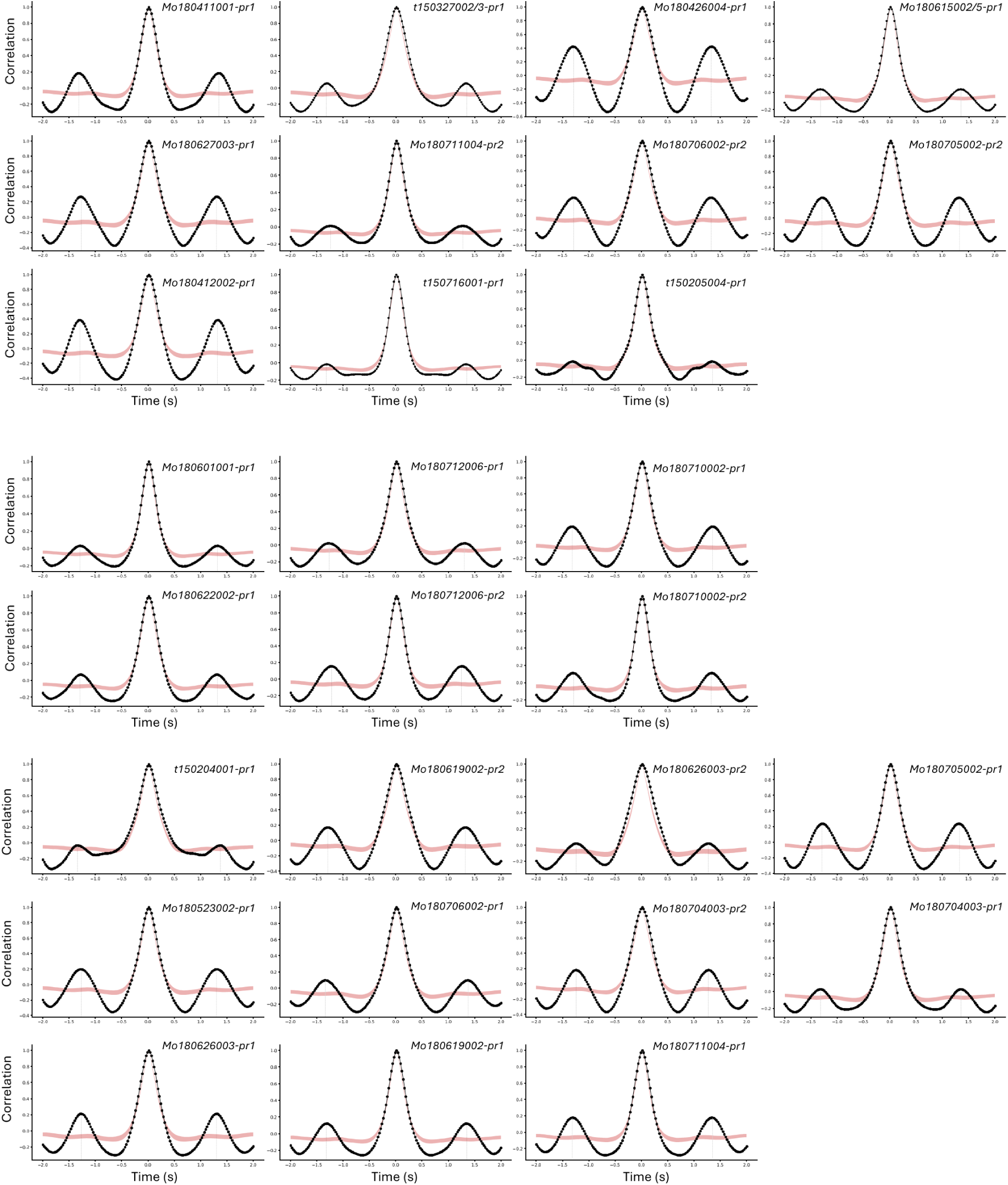
Periodicity of the LD1_color_. Autocorrelation of the cosine similarity structure of LD_1_ (color) across time, shown for all sites exhibiting significant periodicity. The correlation value (black curve) was estimated by averaging all diagonal autocorrelations (lag τ ± 2s) after SC1. Shaded red areas indicate the 5^th^-95^th^ percentile range obtained from block-shuffled surrogate data (block size 0.6-1s). Significant non-zero peaks indicate the estimated period.

**Suppl.Figure 9.**
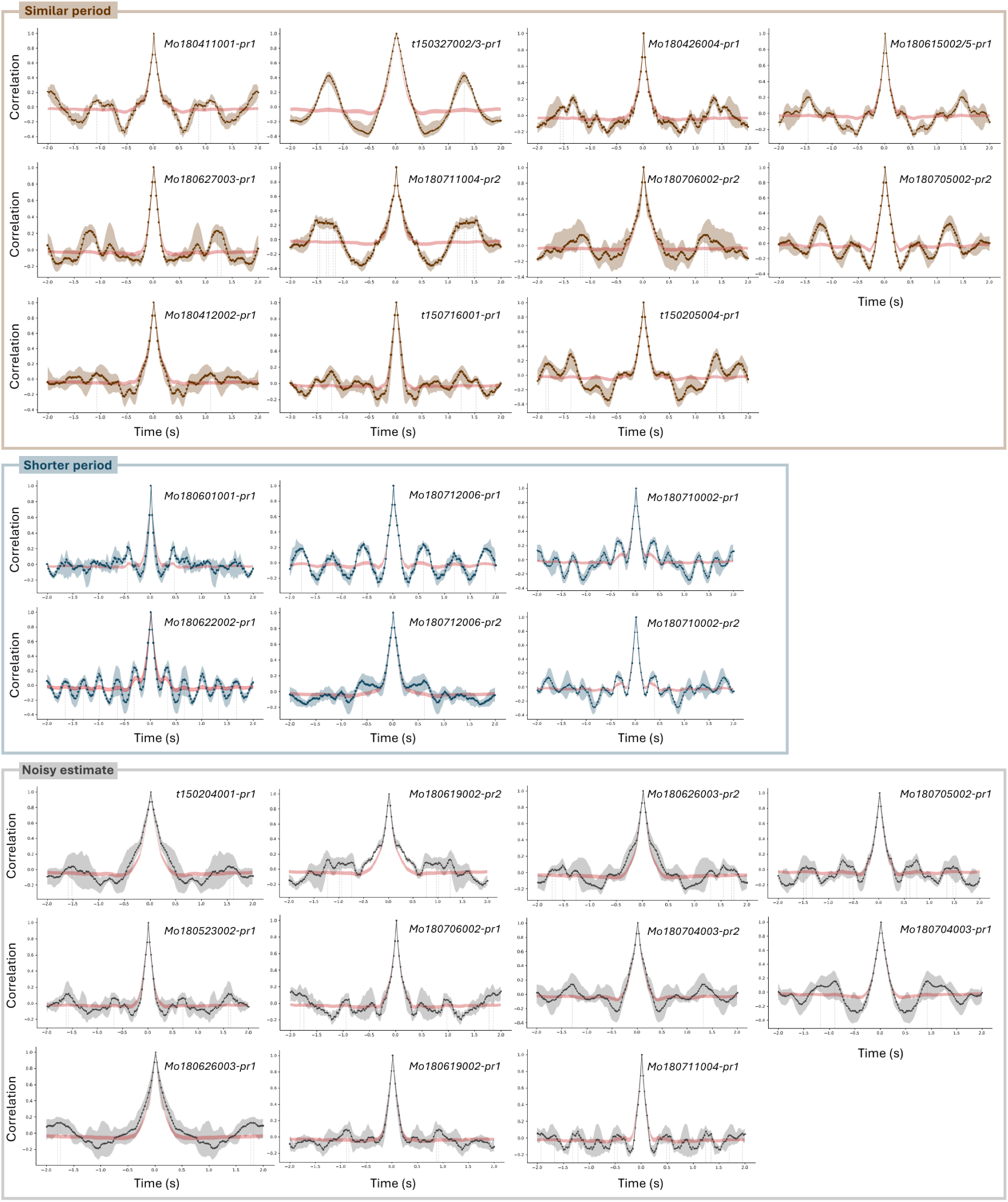
Periodicity of the layer index from LD1_color_. Autocorrelation of the layer index from LD1 (color). The bootstrapped mean autocorrelation is displayed as a dotted colored line, color-coded for the kind of periodicity displayed. Shaded colored areas correspond to the 5^th^-95^th^ confidence interval of the bootstraps. Shaded red areas indicate the 5^th^-95^th^ percentile range obtained from block-shuffled surrogate data (block size 0.3-0.8s). Dashed vertical lines mark detected peaks.

## Notes

### Competing Interest Statement

The authors have declared no competing interest.

### Summary of Updates

Last figure revised; Supplementary figures added; Last results section and discussion updated accordingly.

